# Inference of the worldwide invasion routes of the pinewood nematode *Bursaphelenchus xylophilus* using approximate Bayesian computation analysis

**DOI:** 10.1101/452326

**Authors:** Sophie Mallez, Chantal Castagnone, Eric Lombaert, Philippe Castagnone-Sereno, Thomas Guillemaud

## Abstract

Population genetics have been greatly beneficial to improve knowledge about biological invasions. Model-based genetic inference methods, such as approximate Bayesian computation (ABC), have brought this improvement to a higher level and are now essential tools to decipher the invasion routes of any invasive species. In this paper, we performed ABC random forest analyses to shed light on the pinewood nematode (PWN) worldwide invasion routes and to identify the source of European populations. Originating from North America, this microscopic worm has been invading Asia since 1905 and Europe since 1999, causing tremendous damage on pine forests. Using microsatellite data, we demonstrated the existence of multiple introduction events in Japan (at least two involving individuals originating from the USA) and China (one involving individuals originating from the USA and one involving individuals originating from Japan). We also found that Portuguese samples had a Japanese origin. We observed some discrepancies between descriptive genetic methods and the ABC method, which are worth investigating and are discussed here. The ABC method helped clarify the worldwide history of the PWN invasion, even though the results still need to be considered with some caution because the features of the PWN and the genetic markers used probably push the ABC method to its very limits.

## Introduction

Biological invasions are recognized as one of the main threats to biodiversity (Walker & Steffen, 1997) and are fully integrated in global environmental changes induced by humans (Ricciardi, 2007; Sala et al., 2000; Vitousek, Dantonio, Loope, & Westbrooks, 1996; Wilcove, Rothstein, Dubow, Phillips, & Losos, 1998). Owing to their mainly irreversible aspect (Mooney & Cleland, 2001; but see Simberloff, 2009, for examples of successful eradications) and as their number is growing, studying them constitutes huge challenges at ecological, economical, societal and scientific scales (Mack et al., 2000).

Understanding biological invasions necessitates identifying the history of the invasive process, including routes of invasion. The routes of invasion are defined as the pathways followed by organisms (individuals, seeds…) between their source populations and the invasive populations they formed in a new area. Their identification is a crucial step, whose accuracy is improved by the use of genetic data and analyses (Estoup & Guillemaud, 2010).

Several studies of many invasive species have shown the usefulness of genetic data and analyses to decipher the invasion routes (Boucher et al., 2013; Ciosi et al., 2008; Facon et al., 2003; Fontaine, Gladieux, Hood, & Giraud, 2013; Kelager, Pedersen, & Bruun, 2013; Papura et al., 2012; Perdereau et al., 2013; Rollins, Woolnough, Wilton, Sinclair, & Sherwin, 2009; Wan, Liu, & Zhang, 2012). Recently, quantitative methods were proposed, among which the approximate Bayesian computation (ABC) method (Beaumont, Zhang, & Balding, 2002). ABC relies on the simulation of genetic data following different demographic scenarios defined by the user, and presents several advantages compared to descriptive methods, including (i) the statistical comparison of different invasion scenarios, (ii) the integration of historical and biological knowledge and (iii) the evaluation of the ability to select the true scenario. In brief, various models supposed to explain the data are compared based on the similarity between summary statistics simulated from these models and from prior distributions of historical and genetic parameters on one hand and summary statistics computed from the actual samples on the other hand. This tool has helped reconstructing invasion histories of multiple species (Ascunce et al., 2011; Barres et al., 2012; Boissin et al., 2012; Fraimout et al., 2017; Guillemaud et al., 2015; Lombaert et al., 2010a; Miller et al., 2005; Pascual et al., 2007; Rius, Turon, Ordonez, & Pascual, 2012; Sherpa et al., 2019).

The aim of the present study was to decipher the invasion history of the pinewood nematode (PWN), *Bursaphelenchus xylophilus* (Steiner & Buhrer, 1934; Nickle, 1970; Nematoda: Aphelenchoididae). The PWN is a microscopic worm, exclusively sexual (Futai, 2013), responsible for the pine wilt disease (Mamiya, 1972, 1976, 1983) which annually kills millions of pine trees worldwide (Mamiya, 1988; Soliman et al., 2012; Suzuki, 2002; Vicente, Espada, Vieira, & Mota, 2011). The PWN is native to North America, namely Canada and the USA (Dropkin et al., 1981; Kiritani & Morimoto, 2004). The first invasive outbreak of the PWN was observed in Japan in 1905. The species was then observed in China in 1982, in Taiwan before 1985 and in South Korea in 1988 (Futai, 2013; Mamiya, 1988; Moon, Cheon, & Lee, 2007). In 1999, the PWN was observed in Europe, first in Portugal, and then Spain and Madeira Island in 2008 and 2009 respectively (Abelleira, Picoaga, Mansilla, & Aguin, 2011; Fonseca et al., 2012; Mota et al., 1999). To date, the origin of PWN European populations is still unclear. Some authors have proposed an Asian origin for European invasive populations of PWN (Figueiredo et al., 2013; Valadas, Barbosa, Espada, Oliveira, & Mota, 2012). However, descriptive genetic methods based on genetic distances and clustering analyses recently failed to distinguish between an American and an Asian origin for European outbreaks (Mallez et al., 2015). Analyses based on *F*_*ST*_ values (Weir & Cockerham, 1984) and mean individual assignment likelihoods (Paetkau, Slade, Burden, & Estoup, 2004) suggested an American origin for all Portuguese samples, while analyses based on Cavalli-Sforza and Edwards’ distances (Cavalli-Sforza & Edwards, 1967) and Bayesian clustering (Pritchard, Stephens, & Donnelly, 2000) suggested a Japanese origin for these samples. In this study, we complemented the 34 population samples (770 individuals) used by Mallez et al. with 14 new ones (310 individuals), and used ABC to clarify the invasion history of the PWN (Miller et al., 2005; Pascual et al., 2007).

## Methods

### Sampling and genotyping

In this study, 48 site samples (among which 34 already used in Mallez et al., 2015), representing a total of 1080 individuals were analyzed: 28 site samples from the USA (554 individuals) representing the native area, with a strong focus on Nebraska state (14 site samples, 286 individuals), and 7 site samples from Japan (210 individuals), 9 from Portugal/Madeira (169 individuals) and 4 from China (147 individuals) representing the invaded areas. The samples are listed and described in Table 1, and their geographical locations are shown in Figure S1. No permission was required to collect samples of PWN from the infested areas and we obtained an official agreement from the French authorities (#2012060-0004) for the importation and manipulation of this quarantine organism at Institut Sophia Agrobiotech facilities. All the individuals were extracted from wood samples collected directly in the field, one tree per location, using a sieve or the Baermann method (Viglierchio & Schmitt, 1983), and each site sample consisted of seven to 41 individuals (without separation according to sex or developmental stage). All samplings were performed in 2012, except for China where they were carried out in 2013. Specimens were washed several times in distilled water and stored in DESS (Yoder et al., 2006) at 4°C before DNA was extracted. The historical knowledge about the invasion of the PWN and its spread within each invaded area (Abelleira et al., 2011; Fonseca et al., 2012; Futai, 2013; Mamiya, 1988; Moon et al., 2007; Mota et al., 1999) allowed us to associate a date of first observation to the different invasive outbreaks (Table 1).

**Table 1:**
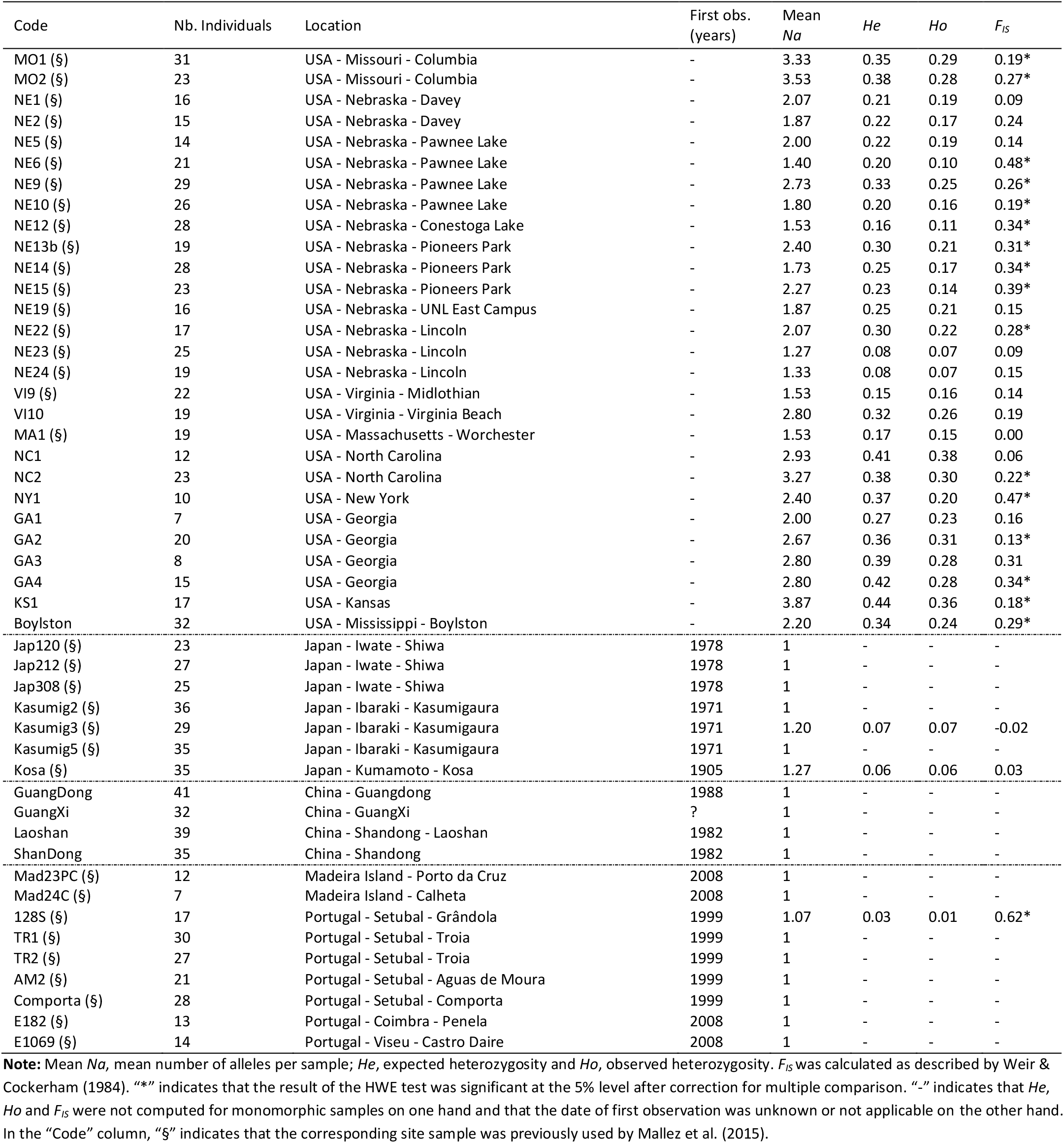
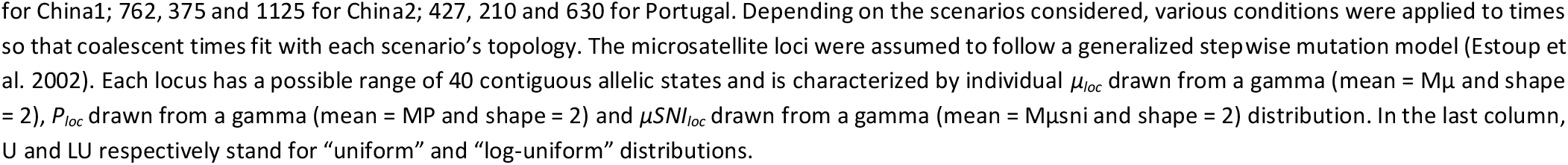
Characteristics and population genetics summary statistics of the 48 site samples of the pinewood nematode used in this study.

The DNA extraction of each single individual was carried out by thermal shock (Castagnone, Abad, & Castagnone-Sereno, 2005) and 16 microsatellite loci were then amplified as described by Mallez *et al*. (2015; 2013). The locus M26 was excluded due to a lack of amplification for one Chinese site sample. The analyses presented here were thus conducted with 15 microsatellite loci.

### Descriptive genetic analyses

For each site sample, we computed the mean number of alleles (Mean *Na*) and the observed (*Ho*) and expected (*He*) heterozygosities per site sample with Genetix version 4.05 (Belkhir, Borsa, Chikhi, Raufaste, & Bonhomme, 1996-2004). We tested each site sample for a deviation from Hardy-Weinberg equilibrium (HWE) with Genepop version 4.1.3 (Rousset, 2008) and we quantified the inferred deviations to HWE by computing the *F*_*IS*_ estimate of Weir and Cockerham (1984) with Fstat version 2.9.3.2 (Goudet, 2002). Linkage disequilibria (LD) between all pairs of loci were also tested with Genepop (Rousset, 2008). The significance level of multiple tests of HWE and LD were adjusted by applying the false discovery rate (Benjamini & Hochberg, 1995) and sequential Bonferroni (Sokal & Rohlf, 1995) corrections, respectively.

### Evolutionary relationships between the different site samples

We used descriptive genetic methods to (i) determine the level of genetic structure existing in each area under study, (ii) reduce the data and select some site samples to include in the ABC analyses (see below) and (iii) guide the choice of the invasive scenarios to compare in these latter analyses.

First, we evaluated the level of genetic differentiation between all site sample pairs with Genepop (Rousset, 2008). Non independent multiple testing implied to adjust the significance level with the sequential Bonferroni method (Sokal & Rohlf, 1995). To assess differentiation between site samples, pairwise *F*_ST_ (Weir & Cockerham, 1984) corrected for null alleles (observed for some of the site samples and loci, see Mallez et al., 2015) were computed with FreeNA (Chapuis & Estoup, 2007). As *F*_ST_ is dependent on the within-population variability (Charlesworth 1998; Jost 2008; Gerlach et al. 2010), we also calculated the pairwise *D*_*EST*_ of Jost (2008) with the SMOGD software (Crawford, 2010).

Second, we performed Bayesian clustering analyses within each geographical area with Structure version 2.3 (Pritchard, Stephens, & Donnelly, 2000). These analyses were carried out only for samples within the USA and China given that the other areas (i. e., Japan and Portugal) had been analyzed previously (Mallez et al., 2015). We used the admixture model with correlated allele frequencies. The number of clusters, *K*, varied from 1 to the total number of site samples per area, i.e., 18 for the USA and 4 for China. Twenty independent runs per *K* were carried out with 10^6^ MCMC iterations each, following a burn-in period of 2×10^5^ iterations. For each Structure analysis, the Clumpak server (Kopelman, Mayzel, Jakobsson, Rosenberg, & Mayrose, 2015), which combines the Clumpp (Jakobsson & Rosenberg, 2007) and Distruct softwares (Rosenberg, 2004), was used to postprocess the Structure outputs, i.e., (i) to determine the highest level of genetic structure using the *ΔK* of Evanno (2005) and (ii) to identify the most frequent clustering pattern for each value of *K* (across the 20 independent runs) and to display the corresponding bar plots.

Third, the Structure software was also run with all site samples from all areas as previously described with *K* varying from 1 to 10 (which appeared to be high enough to capture the genetic pattern existing between the native and the invaded areas, see Results). For this Structure analysis, the Clumpak server (Kopelman et al., 2015) was also used to postprocess the Structure outputs as presented above. Examination of Structure results allowed us to propose putative historical relationships existing between the different site samples.

Finally, a neighbor-joining tree based on Cavalli-Sforza & Edwards’ distances (Cavalli-Sforza & Edwards, 1967), corrected for null alleles with FreeNA (Chapuis & Estoup, 2007) was plotted and the mean individual assignment likelihoods (denoted L_*i*→*s*_, Paetkau, Slade, Burden, & Estoup, 2004) of each invasive site sample *i* to each putative source sample *s* were computed with GeneClass2 version 2.0 (Piry et al., 2004).

### ABC analyses

All ABC model-choice analyses were performed with DIYABC Random Forest v1.0 (Collin et al. 2021). This program implements in the context of ABC a supervised machine-learning algorithm called the Random Forest (Breiman, 2001). Briefly, this non-parametric classification method uses hundreds of bootstrapped decision trees (creating the so-called forest) to perform classification using a set of predictor variables, here the summary statistics.

To define the scenarios to compare, we took into account the historical knowledge (i.e., dates of first observation), results from this study and previous analyses (Mallez et al., 2015). Particular site samples representing genetic units inferred from Structure analyses were used for the ABC analyses. As proposed by Lombaert *et al*. (2014), we performed a step-by-step ABC analysis: we first tried to clarify the Asian invasion history (i.e., the oldest invasive area) and then integrated Europe (i.e., the most recent invasive area) into the scenarios. Thus, we first focused on establishing the number of independent introduction events in Japan (step 1) and in China (step 2), separately. Then, we studied the relationships between Japan and China (steps 3 and 4). Finally, we integrated the Portuguese site sample into the inferred Asian invasion scenario to determine its origin among all possible sources (steps 5 and 6). See Results for details, and Figure S2 for a graphical representation of a few competing scenarios. All configuration files are available (Mallez et al. 2021) at https://doi.org/10.5281/zenodo.4681180.

Within ABC analysis, the number of simulated datasets for each scenario was not uniform, because we took into account the dates of first observations and their prior range. For example, a Chinese population could be the source of a Japanese population in one scenario, but the opposite was more likely *a priori*. For each analysis, we simulated a mean of 20,000 datasets per competing scenario. To reduce the datasets, we used all available summary statistics (SuSts): the mean number of alleles, the mean expected heterozygosity and the mean allelic size variance per population and pairs of populations, the ratio of the number of alleles on the allelic size range (Garza & Williamson, 2001), the mean individual assignment likelihoods of population *i* to population *j* (Pascual et al., 2007), the pairwise *F*_ST_ (Weir & Cockerham, 1984), the shared allele (Chakraborty & Jin, 1993) and *dµ*^2^ (Goldstein, Linares, Cavalli-sforza, & Feldman, 1995) distances between pairs of populations, the maximum likelihood estimate of admixture proportion for each trio of populations (Choisy et al. 2004) and the axes obtained from a linear discriminant analysis on summary statistics (Estoup et al. 2012). We grew a classification forest of 1,000 trees based on all simulated datasets. The random forest computation applied to the observed dataset provides a classification vote representing the number of times a model is selected among the 1000 trees. The scenario with the highest classification vote was selected as the most likely scenario. We then estimated its posterior probability by way of a second random forest procedure of 1,000 trees as described by Pudlo et al. (2016).

To evaluate the robustness of our inferences, we performed the ABC analyses with two different sets of prior distributions (hereafter referred to as “main” and “alternative” prior sets; Table 2). The generation time of the PWN *in natura* is not well-known due to a lack of direct observations of the PWN inside the tree. Considering that the species reproduces from June to September and estimating an average summer temperature of 25°C where it is currently distributed, thirty generations per year was the most realistic, and this value was thus used for all other ABC analyses. However, we tested the effect of choosing other generation times, 15 and 45 generations per year, with the main prior set. We also evaluate the global performance of our ABC Random Forest scenario choice, we computed the prior error rate based on the available out-of-bag simulations (i.e. simulations that are not used in tree building at each bootstrap). The prior error rate corresponds to the proportion of ABC analyses that do not give the highest probability to the true scenario, and thus informs us on our ability to select the true scenario. Finally, we used DIYABC 2.1.0 (Cornuet et al. 2014) to check the adequacy between the observed dataset and the selected scenario, i.e., the capacity of this scenario to generate datasets similar to the observed one (model checking analysis, Cornuet et al. 2010). To do so, we simulated a new reference table consisting of 100,000 datasets of the selected scenario. Then, 1,000 new datasets were simulated by drawing the parameter values from the posterior distributions of the selected scenario. Each observed SuSt was confronted to the distribution formed by the SuSts obtained from the new simulated datasets, which gave a rejection probability for each observed SuSt. A large number of probabilities being obtained, we adjusted the significance threshold with the false discovery rate method (Benjamini & Hochberg, 1995).

**Table 2:**
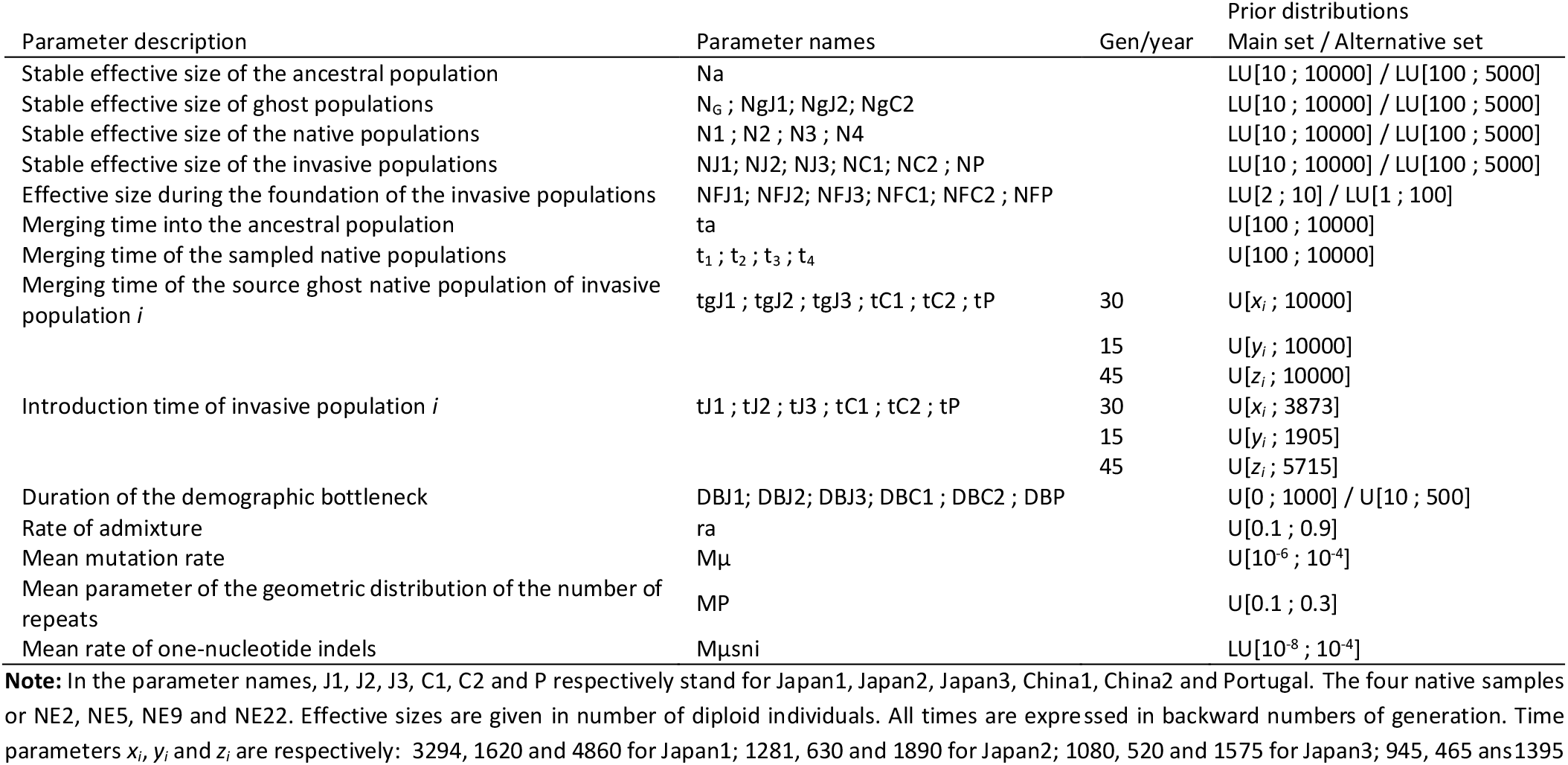
Definition and prior distributions of parameters used in the scenarios of invasion considered in the ABC analyses.

## Results

### Descriptive genetic analyses

The genetic features of the site samples are shown in Table 1. The American site samples displayed a genetic diversity low to moderate with mean *Na* ranging from 1.27 to 3.87 and *He* from 0.08 to 0.44. More than half of the site samples were not at HWE and significant LD were observed for 13 tests out of the 1,084 realized, after correction. This may be due to the presence of null alleles, as already observed by Mallez *et al*. (2015). Raw data are publicly available (Mallez et al. 2021).

All site samples from the invaded areas exhibited very low levels of genetic diversity owing to the presence of numerous monomorphic loci: 9 for Japan, 10 for China and 14 for Portugal/Madeira out of the 15 loci under study. In Japan, three site samples (Jap212, Jap308 and Jap120) appeared completely undifferentiated, with all the individuals sharing the same multi-locus genotype. In Portugal/Madeira, 162 individuals out of the 169 analyzed had the same multi-locus genotype and only one site sample, 128S, was different from the others. In China, all loci were monomorphic within site samples and three site samples out of the four studied presented the same multi-locus genotype for all their individuals.

### Relationships between site samples within geographical areas

The analysis of the American site samples led to the general observation of a strong genetic structure among PWN populations in USA, in good agreement with data from a previous sampling in the same area (Mallez et al., 2015). Indeed, almost all the American site samples appeared significantly differentiated after correction for multiple tests (Fisher method of probability combination on loci, *p* < 10^−2^). Only one test, between NC2 and GA3, was not significant (Fisher method, *p* = 0.11). *F*_*ST*_, corrected for null alleles, reached high values and ranged from 0.036 to 0.76 (Table S1) and *D*_*EST*_ ranged from 0.0041 to 0.329 (Table S1). This strong genetic differentiation pattern was supported by the Structure analysis, which revealed biological relevant clustering of individuals within site samples for high values of *K*, although the *ΔK* inferred *K* = 2 as the main structure (Figure S3). Different clustering patterns were identified with Clumpp, but they exhibited similar results, i.e., a strong genetic structure (Figure S3). In Japan, all site samples were also significantly differentiated, except the three undifferentiated ones mentioned above (Fisher method, *p* < 10^−5^) with *F*_*ST*_ corrected for null alleles ranging from 0.627 to 0.995 and *D*_*EST*_ ranging from 0.0042 to 0.0938 (Table S1). In China, all site samples were undifferentiated except Guangdong that appeared significantly differentiated from the others (Fisher method, *p* < 10^−5^) with a high mean corrected *F*_*ST*_ equal to 0.997 and a lower mean *D*_*EST*_ equal to 0.106 (Table S1). This result was confirmed by the Bayesian clustering analysis, which inferred a relevant clustering of individuals for *K* = 2, Guangdong being pulled apart from the others Chinese site samples (Figure S4). In Portugal/Madeira, all site samples were undifferentiated except the site sample 128S, that was significantly different from the others (Fisher method, *p* < 10^−5^) with a mean corrected *F*_*ST*_ equal to 0.341 and a mean *D*_*EST*_ equal to 0.0005 (Table S1). For samples from Japan and Portugal/Madeira, see also Mallez et al (2015; Figure S5) for additional information. In view of the presence of undifferentiated site samples within Japan, China and Portugal/Madeira, only one of these site samples was conserved in each invaded area for simplification. The conserved site samples were the following: Jap212, Laoshan, TR1 and Mad23PC for Japan, China, Portugal and Madeira, respectively.

### Relationships between site samples among geographical areas

All the site samples from different invaded areas appeared significantly differentiated between them and from the American site samples (Fisher method, *p* < 10^−5^). NE2 was the American site sample with the lowest differentiation from site samples from invaded areas based on the *D*_*EST*_ (mean value = 0.034). Another American site sample, NE9, was selected based on the corrected *F*_*ST*_ (mean value = 0.469), NE2 being the second lowest one with this measure (mean value = 0.541). The neighbor-joining tree confirmed that the site sample NE2 is the closest site sample to all site samples from the invaded areas and that the Portuguese site samples are closer to a Japanese site sample than a Chinese or an American site samples (Figure 1).

**Figure 1:**
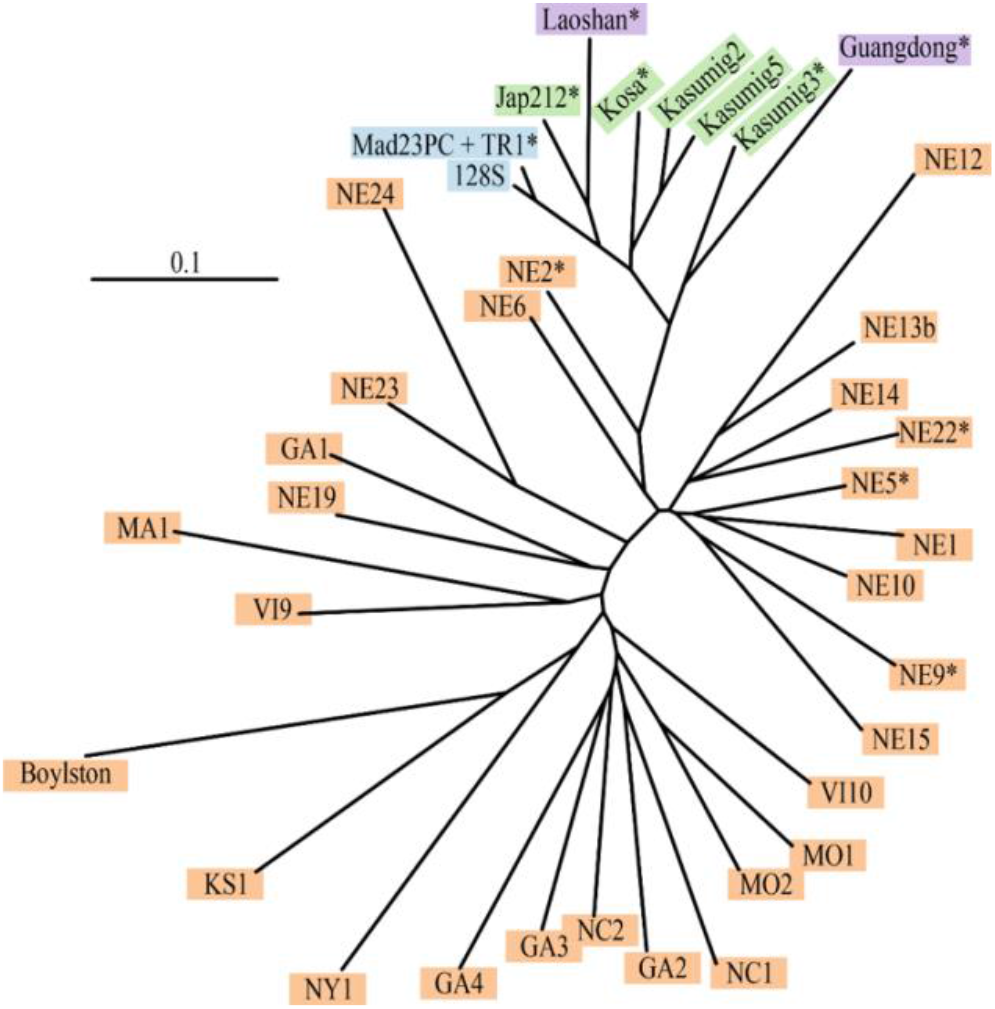
Neighbor-Joining tree built with all the pinewood nematode samples used in this study. Tree based on Cavalli-Sforza and Edwards distances (Cavalli-Sforza and Edwards 1967), corrected for null alleles (Chapuis and Estoup, 2007). The American (native) samples were highlighted in orange, the Japanese samples in green, the Chinese samples in purple and the Portuguese samples in blue. “*” indicates that the corresponding site sample was used in the ABC analyses.

For the Bayesian clustering analysis, the Δ*K* inferred *K*=2 as the main structure (Figure S6A), in which all the invasive site samples pulled apart from the native American site samples (Figure 2). From *K*=3, different clustering patterns were observed (Figures 2 and S6B), in congruence with the sub-structure existing within each of the invaded and native areas. However, from *K*=8, the clustering pattern obtained within the invasive area appeared consistent between runs and identified four groups: one group with Mad23PC, 128S, TR1, Jap212 and Laoshan; another one with Kasumig2, Kasumig5 and Kosa; and Kasumig3 and Guangdong, that clustered separately from one another (Figure 2). Finally, almost all the site samples from the invaded areas were assigned to NE2 with the highest mean individual assignment likelihood (Figure S7). Only Guangdong and Laoshan were assigned to different samples, NE5 and Kosa, respectively (Figure S7).

**Figure 2:**
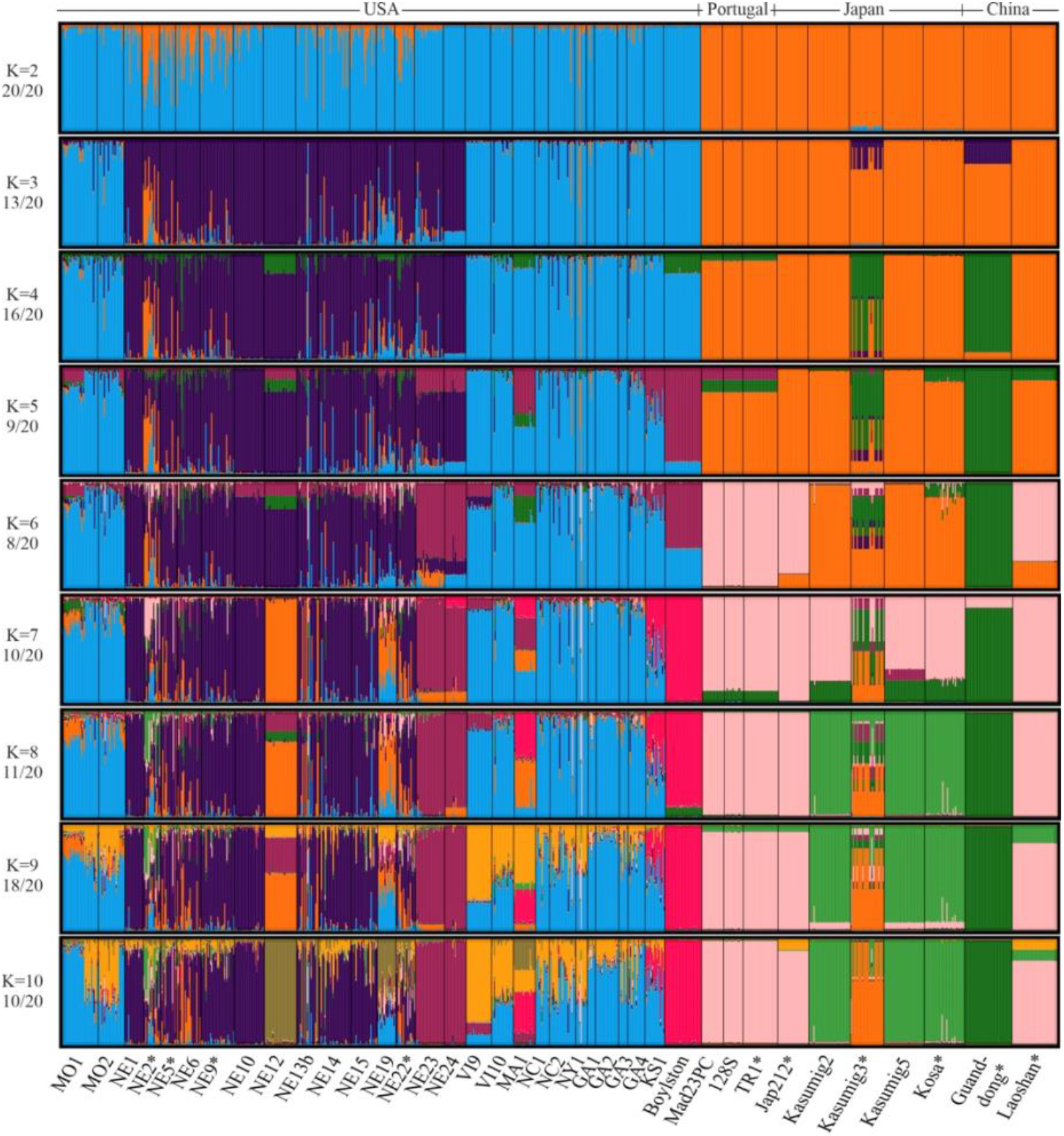
Genetic structure of the pinewood nematode samples used in this study. Bar plots of the coefficients of co-ancestry obtained in STRUCTURE analyses with several values of *K*. Each bar corresponds to one individual nematode and each cluster is represented with a particular color. The major clustering patterns are shown (see Figure S6B for the minor ones, when applicable). “*” indicates that the corresponding site sample was used in the ABC analyses.

Summarizing and considering the results presented above, the native site sample NE2 was the closest to all invasive site samples but Guangdong. Thus, it was used to represent the native area in the competing scenarios of the ABC analyses, together with three other genetically closed samples (NE5, NE9 and NE22). In view of the incomplete sampling of the native area (no sample from Canada was obtained) and the strong genetic structuration observed in the USA, it is unlikely that one of our native sample is representative of the true source population. An unsampled “ghost” population diverging from the group of American samples was thus included in all scenarios (Figure S2). We also considered the following relevant genetic units to characterize the invaded areas in the following ABC analyses:

– Japanese genetic unit 1, hereafter called Japan1, represented by Kosa, the closest sample to the introduction point in Japan and considered coming from the native area (Japan being the first known outbreak area of the PWN);
– Japanese genetic unit 2, hereafter called Japan2, represented by Kasumig3;
– Japanese genetic unit 3, hereafter called Japan3, represented by Jap212;
– Chinese genetic unit 1, hereafter called China1, represented by Laoshan;
– Chinese genetic unit 2, hereafter called China2, represented by Guangdong;
– Portuguese genetic unit, hereafter called Portugal, represented by TR1.

### ABC analyses

The results of the first step indicated that at least two native introductions occurred in Japan, with Japan1 being independent from Japan2, and Japan1 being the source of Japan3, with posterior probability of 0.774 (Table 3, Table S2, Figure S2A). A similar result was observed in China, with the second step: China1 was independent from China2 with posterior probability of 0.963, suggesting two introduction events in China (Table 3, Table S2). The third step’s ABC analysis, carried out between site samples from Japan and China, revealed that Japan1 was probably the source of China1 with posterior probability of 0.573 (Table 3, Table S2, Figure S2B). This result was then confirmed in the fourth step in which we tested, on the basis of the Structure patterns, the hypothesis that Japan2 may have derived from an admixture between China2 on the one hand, and either the native area or Japan1 on the other hand, with posterior probability of 0.567 for the scenario without admixture (Table 3, Table S2). In the fifth step, involving all invaded areas, we found that Japan1 was likely the source of the Portuguese population, with posterior probability of 0.682 (Table 3, Table S2, Figure 3A). The same final scenario (Figures 3A and S2C) was selected in the sixth step, with posterior probability of 0.567, when it was compared to (i) a “unique invasive bridgehead” scenario (Lombaert et al., 2010b), in which an unsampled “ghost” invasive population, originating from the native area, was the source for Japan1 (being itself the source for Japan3, China1 and Portugal), Japan2 and China2, and (ii) to a “unique native origin” scenario, in which Japan1, Japan2 and China2 all derived from the exact same native “ghost” population (Table 3, Table S2).

**Table 3:** Results of the ABC analyses carried out to infer the history of the pinewood nematode invasion. Prior error rates (PER) and posterior probabilities (post-prob) of the selected scenarios are given for the main prior set, assuming 30 generations per year. Full results for all scenarios, and all prior sets and time scales are shown in Table S2.

| Step<br>(nb SuSts) | Nb<br>scenarios | PER | Selected scenario | Post-prob |
| --- | --- | --- | --- | --- |
| **1: Japan**<br>(301) | 8 | 0.273 | S4: Nat→Japan1→Japan3 ; Nat→Japan2 | 0.774 |
| **2: China**<br>(204) | 3 | 0.171 | S1: Native→China1 ; Native→China2 | 0.963 |
| **3: Japan/China**<br>(576) | 24 | 0.338 | S2: Nat→Japan1→Japan3 & China1 ; Nat→Japan2 ; Nat→China2 | 0.573 |
| **4: Japan/China**<br>**admixture** (576) | 3 | 0.311 | S1: Scenario 2 of step 3 | 0.567 |
| **5: Portugal**<br>(760) | 8 | 0.208 | S2: Nat→Japan1→Japan3 & China1 & Port ; Nat→Japan2 ; Nat→China2 | 0.682 |
| **6: Invasive**<br>**bridgehead** (760) | 3 | 0.356 | S1: Scenario 2 of step 5 | 0.567 |

**Note:** “Nat” stands for the native area, represented by a group of 4 samples (NE2, NE5, NE9 and NE22). “Japan1” stands for the Japanese genetic unit 1, represented by the sample Kosa. “Japan2” stands for the Japanese genetic unit 2, represented by the sample Kasumig3. “Japan3” stands for the Japanese genetic unit 2, represented by the sample Jap212. “China1” stands for the Chinese genetic unit 1, represented by the sample Laoshan. “China2” stands for the Chinese genetic unit 2, represented by the sample Guangdong. “Port” stands for the Portuguese genetic unit, represented by the sample TR1.

**Figure 3:**
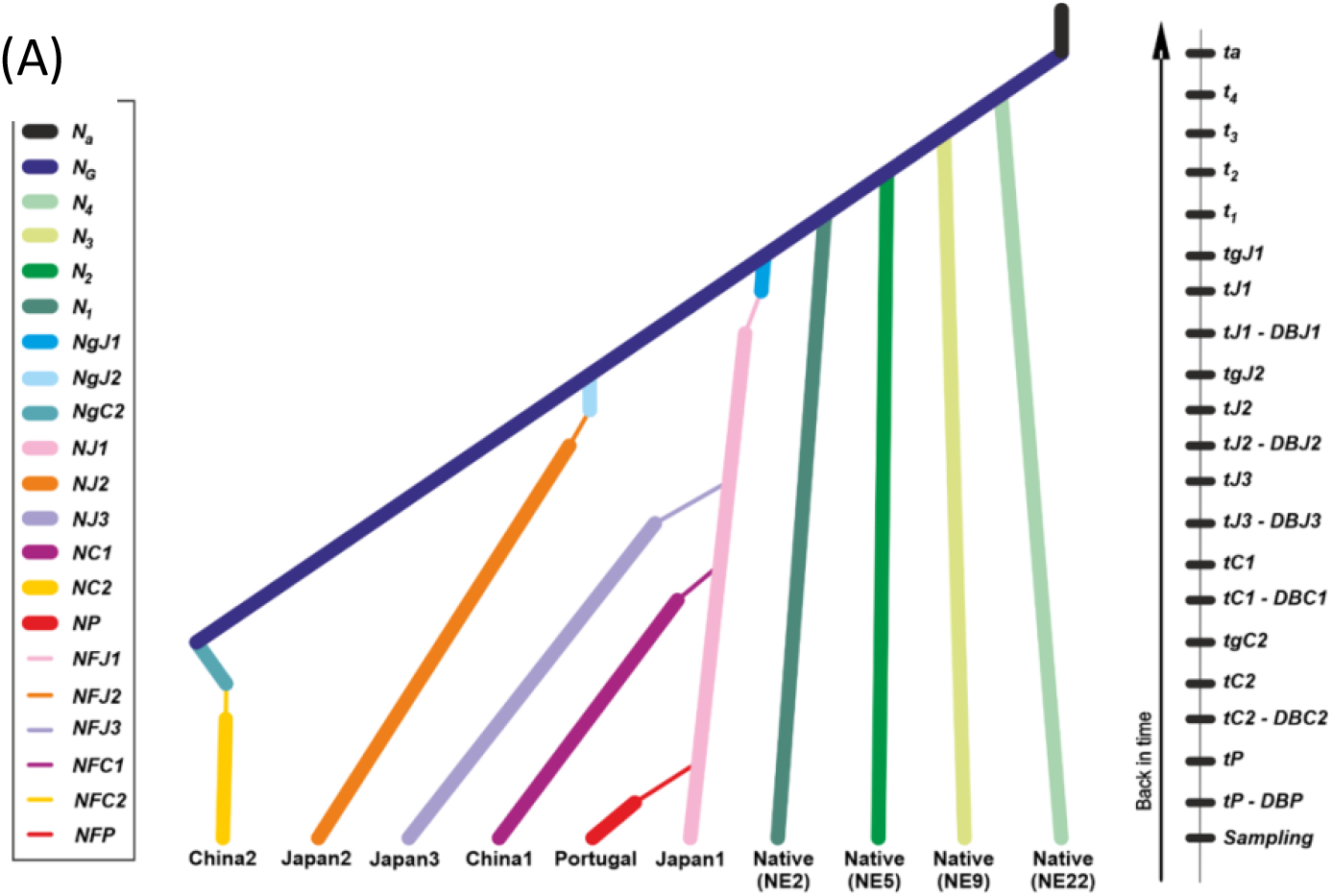
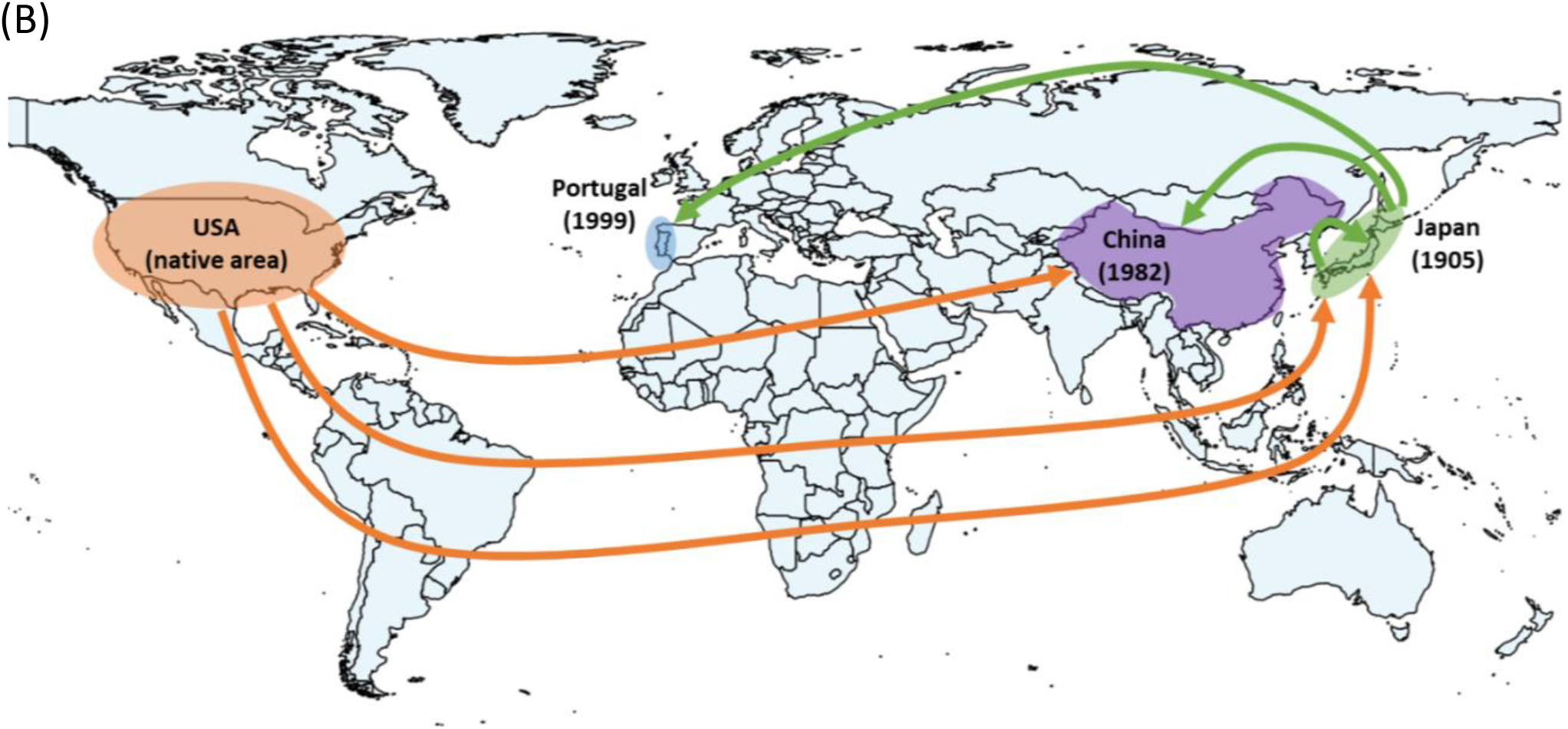
Worldwide invasion routes of the pinewood nematode as inferred from ABC-RF analyses. (A) Graphical illustration of the final selected scenario (step 6, scenario1); (B) Geographic representation of final selected scenario. The year of first infestation for every invaded area is indicated in parenthesis.

For each step, the use of an alternative prior set, a smaller time scale (15 generations/year) or a larger time scale (45 generations/year) essentially confirmed the main results. The exceptions were for steps 1 and 3 for which one analysis out of four provided the largest number of votes for a different scenario (Table S2). The prior error rates appeared moderate for all the analyses (Table 3, Table S2), reflecting a decent, albeit imperfect, ability to distinguish between competing scenarios. Finally, the model checking analysis showed that posterior simulations of the final selected scenario were able to produce data close to the observed ones: none of the 760 observed SuSts were in the extreme 5% tails of the distribution of simulated SuSts after correction by the false discovery rate method, whatever the prior set or time scale used (Table S3).

## Discussion

The main objective of this study was to infer the invasion routes of the PWN using the ABC method. We demonstrated the existence (i) of multiple independent introductions from the native area to Japan and China, and (ii) of a bridgehead population, located in Japan, and which was the secondary source of several populations in Asia and Europe. This inference has not been possible using descriptive population genetics methods performed in a previous study (Mallez et al., 2015). Conversely, the ABC method helped clarify the worldwide history of the PWN invasion, even though the results still need to be considered with some caution. The discrepancies observed between the various methods are worth investigating and are discussed here.

### Invasion routes of the PWN and the origin of Portuguese populations

Despite the low genetic diversity generally present in the PWN populations, we were able to reliably select scenarios (among the ones compared) with good posterior probabilities in most cases and satisfactory quality analyses. The most probable worldwide invasion history of the PWN identified in this study (Figure 3B) includes multiple introductions in Asia and a Japanese origin of the European samples.

The analyses allowed us to reveal the existence of multiple independent events of introduction from the native area to Asia, the USA being the source of two outbreaks in Japan and one in China. Within Asia, the oldest Japanese population has partly spread across the country, but has also acted as a source for at least one population in China. Especially, these results strengthen the hypothesis of two events of introduction in China, the first one near Nanjing in 1982 and the second one near Hong-Kong in 1988, already suggested by modeling approaches in Robinet et al (2009; but see Cheng, Cheng, Xu, & Xie, 2008). Concerning Europe, the ABC analyses of the PWN microsatellites data designated the oldest Japanese population as the most probable source of the Portuguese populations and excluded the native area as a possible source. This result thus confirm what is often presented in the literature based on genetic distance trees (Figueiredo et al., 2013; Fonseca et al., 2012; Metge & Burgermeister, 2008; Valadas et al., 2012; Valadas, Laranjo, Mota, & Oliveira, 2013; Vieira, Burgermeister, Mota, Metge, & Silva, 2007). This result was robust to changes in the parameters’ priors and displayed good posterior model checking results.

This study highlights the central role of the native area in the worldwide history of invasion of the PWN, adding up three as the minimum number of independent introductions from the USA. The other striking result is the bridgehead status of the invasive population represented by our sample of Kosa in Japan, which has a significant role in Asia, but which is also the source of the European invasion. Noteworthy, only one detectable event of introduction seems to have occurred in Portugal. Considering the volume of global trade between Portugal and the worldwide infested areas, this result shows that quarantine and inspection efforts have been globally successful and efficient to date to minimize the number of independent outbreaks and confirms the importance of prevention and pro-active actions to limit new introductions (Simberloff & Rejmanek, 2011). At the European level, ecological and pedo-climatic conditions favorable to the PWN are encountered in many countries (Robinet, Van Opstal, Baker, & Roques, 2011). However, the invasion is currently limited to the Iberian Peninsula (plus Madeira Island). This reinforces the idea that the quarantine regulation measures have been efficient to date to protect Europe from the entry of new invasive PWN populations from abroad, and from a large dissemination of the PWN from the original Portuguese outbreak.

### Different methods with different results

In the case of the PWN invasion, the results obtained from descriptive genetic approaches pointed out different putative origins for the European populations of PWN (Mallez et al., 2015): American based on *F*_*ST*_ values (Weir & Cockerham, 1984) and mean individual assignment likelihoods (Paetkau et al., 2004), and Japanese based on Cavalli-Sforza and Edwards’ distances (Cavalli-Sforza & Edwards, 1967) and Bayesian clustering (Pritchard et al., 2000). The ABC analyses, quantitative and probability-based, allowed us to improve greatly the history of the PWN invasion. Our study highlights discrepancies between the ABC methodology and some of the descriptive approaches, such as the *F*_*ST*_ and the mean assignment likelihoods. It is interesting to notice that these two statistics are among those chosen to summarize the data in the ABC analyses. However, in addition to using other key statistics, the ABC method allows us to consider complex evolutionary scenarios, involving for example intense bottlenecks or unsampled populations. Indeed, previous studies using simulated datasets have shown that *F*_*ST*_ and mean assignment likelihoods, used alone, can be misleading in some situations (Guillemaud et al., 2010; Lombaert et al., 2010; 2018).

On the contrary, the inference of multiple independent introductions from the native area in Asia (Japan1, Japan2 and China2) is not consistent with the Bayesian clustering of individuals (Pritchard et al., 2000) and distance genetic trees (Saitou & Nei, 1987). To our knowledge, such discrepancies have not been observed yet, although a simulation study showed that classic clustering methods such as Structure analyses (Pritchard et al., 2000) may be erroneously interpreted in the context of invasion route inference (Lombaert, Guillemaud, & Deleury, 2018). The origin of the discrepancies we observed is not completely clear to us, but hypotheses may be proposed. They may likely have arisen from the very low level of genetic diversity observed in the PWN populations, which might disturb the analyses of the descriptive genetic methods and thus their outcomes. In an invasion context, a low genetic diversity is usually the translation of strong founder effects, which implies a strong genetic drift. Genetic drift, in turn, is a stochastic process whose consequences are random. Employing methods such as ABC that take these stochastic effects into account may thus be more reliable than other methods that do not, such as Bayesian clustering or distance-based methods (Guillemaud et al., 2010; Estoup & Guillemaud, 2010). In these conditions of low genetic diversity, we can expect descriptive methods to lead to variable results when applied to various realizations of the same historical scenario. We may expect the low level of genetic diversity to produce a genetic structure pattern and/or relationship between site samples that we misinterpret. For instance, if the native area is weakly diversified so that it exhibits a few very frequent alleles, it is probable that two independent introductions from this native area (native → invasive 1 and native → invasive 2) lead to samples closer to each other than to their native area. When analyzed, such results would suggest serial introductions (native → invasive 1 → invasive 2, because invasive1 and 2 appear closer to each other than to native) instead of the true independent introduction scenario, because the interpretation is binary and not based on probability computation. Indeed, the use of simulated datasets enabled Lombaert et al. (2018) to show that multiple introductions from a native population with small effective size, and thus low diversity, could often lead to single introduction patterns with Structure. Conversely, using a model-based stochastic method, such as ABC, a genetic structure with invading samples closer to each other than to the native area may lead to the choice of independent introduction scenario because it is probable to get such structure with this scenario when the diversity is low.

Overall, the ABC method, explicitly based on models, seems appropriate for the PWN which is characterized by a markedly low intra-population diversity and a very strong inter-population genetic structure. However, one should remain cautious when interpreting our results. Indeed, the features of the PWN and of the genetic markers used probably push the ABC method to its very limits. Particularly noteworthy are the sometimes relatively high prior error rates, and the few – albeit rare – scenario selection shifts when using different time scales. Finally, a factor that is probably of particular importance is the incompleteness of our sampling scheme, especially in the native area. The genetic structure is so strong in the latter that the use of ghost populations in our analyses may not be sufficient to capture the full complexity of the true evolutionary history.

Carrying out simulations may help to disentangle the effect of the genetic diversity in the native area on the outcomes of clustering methods and ABC analyses, a work partly performed by Lombaert *et al*. (2018) who did not use ABC analyses. More precisely, analyzing datasets, simulated under chosen scenarios and known levels of genetic diversity, may allow to assess the behavior of these analyses in relation to the level of genetic diversity. These simulations would thus be useful to (i) determine the real impact of the genetic diversity on the outcomes, (ii) assess the expected benefits of increasing the number of genetic markers and (iii) verify that the interpretations made from these methods in the biological invasion framework are appropriate.

## Data accessibility

Complete dataset (microsatellite), STRUCTURE outputs and ABC configuration files were deposited at Zenodo: https://doi.org/10.5281/zenodo.4681180

## Acknowledgements

We would like to thank Pedro Barbosa, Douglas LeDoux, Julia Thompson, Margarida Espada, Paulo Vieira, Jonathan D. Eisenback, Mark Harrell, Manuel Mota, Takuya Aikawa, Mitsuteru Akiba, Hajime Kosaka, Lilin Zhao and Jianghua Sun for technical support, precious help in sampling and fruitful discussions. This work was funded by the EU REPHRAME project (KBBE.2010.1.4-09). Version 6 of this preprint has been peer-reviewed and recommended by Peer Community In Zoology (https://doi.org/10.24072/pci.zool.100006)

## Author Contributions

SM and CC performed microsatellite genotyping. SM performed the analyses, with input from EL. SM, EL, PCS and TG interpreted the results. SM wrote the manuscript, with guidance from PCS and TG, and input from EL. All authors have read and approved the final manuscript.

