## Supplemental file for "Inference of the worldwide invasion routes of the pinewood nematode *Bursaphelenchus xylophilus* using approximate Bayesian computation analysis"

| | |
|---|---|
| *Table S1* | *p. 2* |
| *Table S2* | *p. 4* |
| *Table S3* | *p. 6* |
| *Figure S1* | *p. 7* |
| *Figure S2* | *p. 8* |
| *Figure S3* | *p. 9* |
| *Figure S4* | *p. 12* |
| *Figure S5* | *p. 13* |
| *Figure S6* | *p. 14* |
| *Figure S7* | *p. 15* |

**Table S1:** Measures of differentiation between pairs of samples from native and invasive areas. The $D_{EST}$ of Jost (2008, *Molecular Ecology*, 17:4015-4026), computed using SMOGD (Crawford, 2010, *Molecular Ecology Resources*, 10:556-557), is shown below the diagonal and the $F_{ST}$ of Weir and Cockerham (1984, *Evolution*, 38:1358-1370), corrected for null alleles and obtained with FREENA (Chapuis and Estoup, 2007, *Molecular Biology and Evolution*, 24:621-631), is shown above the diagonal. The color gradients below and above the diagonal are independent.

| | | MO1 | MO2 | NE1 | NE2 | NE5 | NE6 | NE9 | NE10 | NE12 | NE13b | NE14 | NE15 | NE19 | NE22 | NE23 | NE24 | Vi9 | VI10 | MA1 | NC1 | NC2 | NY1 | GA1 | GA2 | GA3 | GA4 |
|---|---|---|---|---|---|---|---|---|---|---|---|---|---|---|---|---|---|---|---|---|---|---|---|---|---|---|---|
| | **MO1** | | 0.065 | 0.294 | 0.246 | 0.279 | 0.275 | 0.226 | 0.328 | 0.388 | 0.176 | 0.221 | 0.314 | 0.193 | 0.250 | 0.415 | 0.500 | 0.334 | 0.133 | 0.429 | 0.127 | 0.112 | 0.260 | 0.208 | 0.137 | 0.104 | 0.086 |
| | **MO2** | 0.022 | | 0.266 | 0.194 | 0.240 | 0.257 | 0.194 | 0.291 | 0.402 | 0.173 | 0.196 | 0.265 | 0.151 | 0.214 | 0.390 | 0.504 | 0.307 | 0.109 | 0.387 | 0.074 | 0.055 | 0.217 | 0.145 | 0.101 | 0.074 | 0.092 |
| | **NE1** | 0.097 | 0.089 | | 0.188 | 0.120 | 0.168 | 0.140 | 0.180 | 0.493 | 0.223 | 0.227 | 0.238 | 0.319 | 0.216 | 0.505 | 0.623 | 0.508 | 0.333 | 0.596 | 0.305 | 0.253 | 0.367 | 0.387 | 0.268 | 0.278 | 0.263 |
| | **NE2** | 0.067 | 0.054 | 0.033 | | 0.145 | 0.161 | 0.108 | 0.250 | 0.500 | 0.236 | 0.188 | 0.247 | 0.311 | 0.110 | 0.448 | 0.641 | 0.474 | 0.301 | 0.553 | 0.238 | 0.221 | 0.355 | 0.330 | 0.260 | 0.272 | 0.257 |
| | **NE5** | 0.100 | 0.079 | 0.009 | 0.014 | | 0.141 | 0.087 | 0.086 | 0.475 | 0.190 | 0.154 | 0.198 | 0.292 | 0.176 | 0.411 | 0.567 | 0.483 | 0.288 | 0.582 | 0.309 | 0.243 | 0.342 | 0.323 | 0.257 | 0.278 | 0.256 |
| | **NE6** | 0.067 | 0.087 | 0.020 | 0.033 | 0.020 | | 0.137 | 0.278 | 0.502 | 0.241 | 0.153 | 0.369 | 0.381 | 0.198 | 0.503 | 0.632 | 0.506 | 0.328 | 0.567 | 0.317 | 0.281 | 0.433 | 0.397 | 0.286 | 0.329 | 0.300 |
| | **NE9** | 0.071 | 0.065 | 0.028 | 0.023 | 0.015 | 0.018 | | 0.153 | 0.411 | 0.116 | 0.108 | 0.167 | 0.290 | 0.121 | 0.430 | 0.501 | 0.398 | 0.251 | 0.482 | 0.206 | 0.215 | 0.317 | 0.296 | 0.233 | 0.241 | 0.202 |
| | **NE10** | 0.116 | 0.095 | 0.016 | 0.038 | 0.004 | 0.031 | 0.023 | | 0.521 | 0.187 | 0.213 | 0.152 | 0.341 | 0.279 | 0.422 | 0.547 | 0.462 | 0.321 | 0.609 | 0.363 | 0.277 | 0.376 | 0.407 | 0.300 | 0.336 | 0.291 |
| | **NE12** | 0.108 | 0.141 | 0.118 | 0.136 | 0.106 | 0.079 | 0.091 | 0.119 | | 0.289 | 0.424 | 0.599 | 0.439 | 0.401 | 0.662 | 0.696 | 0.688 | 0.457 | 0.611 | 0.475 | 0.427 | 0.525 | 0.521 | 0.463 | 0.456 | 0.436 |
| | **NE13b** | 0.036 | 0.061 | 0.055 | 0.052 | 0.045 | 0.035 | 0.030 | 0.037 | 0.049 | | 0.118 | 0.229 | 0.244 | 0.188 | 0.472 | 0.556 | 0.450 | 0.221 | 0.504 | 0.217 | 0.195 | 0.303 | 0.315 | 0.243 | 0.214 | 0.177 |
| | **NE14** | 0.068 | 0.064 | 0.038 | 0.048 | 0.012 | 0.027 | 0.024 | 0.025 | 0.102 | 0.021 | | 0.275 | 0.322 | 0.135 | 0.417 | 0.543 | 0.347 | 0.216 | 0.487 | 0.213 | 0.185 | 0.323 | 0.280 | 0.256 | 0.238 | 0.205 |
| | **NE15** | 0.112 | 0.087 | 0.021 | 0.048 | 0.027 | 0.068 | 0.040 | 0.021 | 0.159 | 0.053 | 0.045 | | 0.357 | 0.294 | 0.574 | 0.613 | 0.456 | 0.269 | 0.602 | 0.284 | 0.227 | 0.323 | 0.383 | 0.271 | 0.286 | 0.256 |
| | **NE19** | 0.068 | 0.036 | 0.064 | 0.080 | 0.058 | 0.096 | 0.089 | 0.074 | 0.107 | 0.093 | 0.094 | 0.059 | | 0.306 | 0.481 | 0.573 | 0.491 | 0.283 | 0.541 | 0.270 | 0.167 | 0.310 | 0.225 | 0.219 | 0.174 | 0.203 |
| **USA** | **NE22** | 0.064 | 0.072 | 0.049 | 0.019 | 0.020 | 0.044 | 0.034 | 0.059 | 0.119 | 0.034 | 0.028 | 0.071 | 0.073 | | 0.476 | 0.611 | 0.441 | 0.285 | 0.510 | 0.217 | 0.205 | 0.311 | 0.297 | 0.268 | 0.247 | 0.230 |
| | **NE23** | 0.120 | 0.106 | 0.048 | 0.052 | 0.033 | 0.062 | 0.080 | 0.044 | 0.132 | 0.095 | 0.060 | 0.088 | 0.069 | 0.091 | | 0.594 | 0.576 | 0.474 | 0.715 | 0.472 | 0.391 | 0.580 | 0.579 | 0.391 | 0.504 | 0.433 |
| | **NE24** | 0.171 | 0.163 | 0.107 | 0.145 | 0.080 | 0.128 | 0.155 | 0.091 | 0.218 | 0.145 | 0.118 | 0.133 | 0.103 | 0.177 | 0.023 | | 0.711 | 0.570 | 0.760 | 0.524 | 0.474 | 0.589 | 0.668 | 0.435 | 0.511 | 0.486 |
| | **Vi9** | 0.109 | 0.095 | 0.099 | 0.105 | 0.087 | 0.119 | 0.123 | 0.069 | 0.286 | 0.122 | 0.048 | 0.085 | 0.122 | 0.113 | 0.096 | 0.151 | | 0.317 | 0.604 | 0.325 | 0.258 | 0.418 | 0.450 | 0.338 | 0.352 | 0.308 |
| | **VI10** | 0.054 | 0.055 | 0.117 | 0.103 | 0.096 | 0.097 | 0.102 | 0.095 | 0.170 | 0.071 | 0.064 | 0.079 | 0.111 | 0.102 | 0.110 | 0.178 | 0.072 | | 0.378 | 0.118 | 0.094 | 0.236 | 0.229 | 0.158 | 0.113 | 0.120 |
| | **MA1** | 0.183 | 0.146 | 0.236 | 0.217 | 0.211 | 0.176 | 0.230 | 0.215 | 0.199 | 0.198 | 0.172 | 0.189 | 0.167 | 0.212 | 0.222 | 0.304 | 0.155 | 0.099 | | 0.405 | 0.379 | 0.474 | 0.541 | 0.418 | 0.442 | 0.441 |
| | **NC1** | 0.064 | 0.035 | 0.118 | 0.099 | 0.133 | 0.120 | 0.103 | 0.135 | 0.216 | 0.068 | 0.069 | 0.092 | 0.112 | 0.072 | 0.152 | 0.200 | 0.080 | 0.042 | 0.149 | | 0.057 | 0.198 | 0.224 | 0.106 | 0.063 | 0.113 |
| | **NC2** | 0.061 | 0.020 | 0.073 | 0.080 | 0.084 | 0.111 | 0.096 | 0.085 | 0.201 | 0.065 | 0.062 | 0.069 | 0.050 | 0.064 | 0.101 | 0.146 | 0.055 | 0.047 | 0.143 | 0.012 | | 0.157 | 0.129 | 0.091 | 0.016 | 0.064 |
| | **NY1** | 0.154 | 0.108 | 0.134 | 0.136 | 0.152 | 0.217 | 0.192 | 0.150 | 0.254 | 0.164 | 0.142 | 0.112 | 0.167 | 0.143 | 0.189 | 0.245 | 0.139 | 0.109 | 0.207 | 0.089 | 0.080 | | 0.248 | 0.245 | 0.125 | 0.165 |
| | **GA1** | 0.090 | 0.048 | 0.097 | 0.090 | 0.078 | 0.112 | 0.118 | 0.122 | 0.115 | 0.128 | 0.088 | 0.114 | 0.051 | 0.099 | 0.122 | 0.177 | 0.093 | 0.092 | 0.168 | 0.110 | 0.051 | 0.094 | | 0.234 | 0.130 | 0.185 |
| | **GA2** | 0.065 | 0.056 | 0.090 | 0.103 | 0.091 | 0.084 | 0.111 | 0.093 | 0.202 | 0.097 | 0.087 | 0.091 | 0.076 | 0.104 | 0.095 | 0.112 | 0.090 | 0.061 | 0.148 | 0.027 | 0.039 | 0.124 | 0.107 | | 0.053 | 0.094 |
| | **GA3** | 0.046 | 0.019 | 0.074 | 0.077 | 0.081 | 0.105 | 0.104 | 0.116 | 0.187 | 0.072 | 0.081 | 0.085 | 0.048 | 0.081 | 0.097 | 0.110 | 0.086 | 0.038 | 0.175 | 0.018 | 0.002 | 0.057 | 0.053 | 0.010 | | 0.035 |
| | **GA4** | 0.054 | 0.049 | 0.088 | 0.094 | 0.094 | 0.115 | 0.088 | 0.093 | 0.192 | 0.057 | 0.073 | 0.100 | 0.091 | 0.071 | 0.126 | 0.162 | 0.083 | 0.062 | 0.212 | 0.076 | 0.032 | 0.112 | 0.094 | 0.057 | 0.009 | |
| | **KS1** | 0.066 | 0.047 | 0.145 | 0.108 | 0.145 | 0.160 | 0.125 | 0.160 | 0.204 | 0.108 | 0.106 | 0.150 | 0.139 | 0.134 | 0.193 | 0.226 | 0.160 | 0.096 | 0.218 | 0.066 | 0.050 | 0.119 | 0.096 | 0.095 | 0.038 | 0.058 |
| | **Boylston** | 0.186 | 0.197 | 0.300 | 0.303 | 0.329 | 0.315 | 0.320 | 0.288 | 0.265 | 0.254 | 0.300 | 0.308 | 0.211 | 0.314 | 0.233 | 0.292 | 0.262 | 0.220 | 0.222 | 0.252 | 0.210 | 0.253 | 0.250 | 0.227 | 0.194 | 0.189 |
| | **Jap308** | 0.135 | 0.135 | 0.044 | 0.041 | 0.060 | 0.079 | 0.078 | 0.102 | 0.217 | 0.159 | 0.094 | 0.096 | 0.096 | 0.063 | 0.107 | 0.213 | 0.121 | 0.173 | 0.270 | 0.141 | 0.110 | 0.166 | 0.116 | 0.130 | 0.102 | 0.132 |
| | **Jap212** | 0.135 | 0.135 | 0.044 | 0.041 | 0.060 | 0.079 | 0.078 | 0.102 | 0.217 | 0.159 | 0.094 | 0.096 | 0.096 | 0.063 | 0.107 | 0.213 | 0.121 | 0.173 | 0.270 | 0.141 | 0.110 | 0.166 | 0.116 | 0.130 | 0.102 | 0.132 |
| | **Jap120** | 0.134 | 0.135 | 0.044 | 0.041 | 0.059 | 0.079 | 0.078 | 0.102 | 0.217 | 0.158 | 0.094 | 0.096 | 0.096 | 0.063 | 0.107 | 0.213 | 0.121 | 0.173 | 0.270 | 0.141 | 0.110 | 0.166 | 0.116 | 0.130 | 0.102 | 0.131 |
| **Japan** | **Kasumig2** | 0.125 | 0.134 | 0.060 | 0.023 | 0.043 | 0.054 | 0.053 | 0.082 | 0.217 | 0.141 | 0.086 | 0.083 | 0.124 | 0.057 | 0.092 | 0.213 | 0.110 | 0.155 | 0.206 | 0.153 | 0.129 | 0.184 | 0.129 | 0.146 | 0.121 | 0.141 |
| | **Kasumig3** | 0.086 | 0.079 | 0.088 | 0.045 | 0.059 | 0.044 | 0.074 | 0.091 | 0.096 | 0.102 | 0.109 | 0.115 | 0.082 | 0.065 | 0.068 | 0.152 | 0.185 | 0.103 | 0.201 | 0.121 | 0.133 | 0.212 | 0.129 | 0.088 | 0.101 | 0.148 |
| | **Kasumig5** | 0.090 | 0.116 | 0.092 | 0.035 | 0.067 | 0.071 | 0.083 | 0.104 | 0.160 | 0.130 | 0.106 | 0.122 | 0.082 | 0.073 | 0.058 | 0.158 | 0.135 | 0.150 | 0.205 | 0.146 | 0.112 | 0.184 | 0.120 | 0.132 | 0.101 | 0.130 |
| | **Kosa** | 0.132 | 0.132 | 0.058 | 0.021 | 0.036 | 0.049 | 0.045 | 0.072 | 0.217 | 0.135 | 0.077 | 0.073 | 0.121 | 0.053 | 0.088 | 0.207 | 0.107 | 0.133 | 0.153 | 0.160 | 0.138 | 0.192 | 0.134 | 0.165 | 0.147 | 0.147 |
| | **GuangDong** | 0.154 | 0.168 | 0.138 | 0.085 | 0.106 | 0.071 | 0.101 | 0.164 | 0.160 | 0.165 | 0.126 | 0.163 | 0.172 | 0.095 | 0.155 | 0.275 | 0.206 | 0.140 | 0.159 | 0.165 | 0.190 | 0.255 | 0.190 | 0.182 | 0.194 | 0.222 |
| **China** | **GuangXi** | 0.153 | 0.176 | 0.094 | 0.050 | 0.072 | 0.087 | 0.068 | 0.107 | 0.227 | 0.147 | 0.123 | 0.110 | 0.149 | 0.090 | 0.155 | 0.279 | 0.178 | 0.227 | 0.340 | 0.216 | 0.179 | 0.257 | 0.165 | 0.192 | 0.179 | 0.154 |
| | **Laoshan** | 0.153 | 0.177 | 0.094 | 0.050 | 0.072 | 0.087 | 0.069 | 0.107 | 0.227 | 0.147 | 0.124 | 0.110 | 0.149 | 0.090 | 0.155 | 0.279 | 0.178 | 0.227 | 0.340 | 0.216 | 0.179 | 0.257 | 0.165 | 0.192 | 0.179 | 0.154 |
| | **ShanDong** | 0.153 | 0.176 | 0.094 | 0.050 | 0.072 | 0.087 | 0.069 | 0.107 | 0.227 | 0.147 | 0.123 | 0.110 | 0.149 | 0.090 | 0.155 | 0.279 | 0.178 | 0.227 | 0.340 | 0.216 | 0.179 | 0.257 | 0.165 | 0.192 | 0.179 | 0.154 |
| | **Mad23PC** | 0.135 | 0.107 | 0.059 | 0.022 | 0.042 | 0.053 | 0.052 | 0.082 | 0.216 | 0.140 | 0.085 | 0.084 | 0.111 | 0.053 | 0.107 | 0.213 | 0.127 | 0.170 | 0.270 | 0.166 | 0.139 | 0.184 | 0.118 | 0.152 | 0.153 | 0.160 |
| | **Mad24C** | 0.134 | 0.105 | 0.058 | 0.022 | 0.041 | 0.052 | 0.051 | 0.081 | 0.216 | 0.139 | 0.084 | 0.084 | 0.110 | 0.052 | 0.107 | 0.212 | 0.126 | 0.169 | 0.270 | 0.164 | 0.138 | 0.182 | 0.118 | 0.151 | 0.151 | 0.157 |
| | **E182** | 0.135 | 0.107 | 0.059 | 0.022 | 0.042 | 0.053 | 0.052 | 0.082 | 0.216 | 0.140 | 0.085 | 0.085 | 0.111 | 0.054 | 0.107 | 0.213 | 0.127 | 0.170 | 0.270 | 0.166 | 0.140 | 0.184 | 0.118 | 0.153 | 0.153 | 0.160 |
| | **128S** | 0.135 | 0.104 | 0.059 | 0.022 | 0.042 | 0.053 | 0.052 | 0.082 | 0.217 | 0.140 | 0.085 | 0.085 | 0.112 | 0.054 | 0.107 | 0.213 | 0.116 | 0.168 | 0.270 | 0.159 | 0.130 | 0.184 | 0.116 | 0.149 | 0.149 | 0.154 |
| **Portugal** | **E1069** | 0.135 | 0.107 | 0.059 | 0.022 | 0.042 | 0.053 | 0.052 | 0.082 | 0.216 | 0.140 | 0.085 | 0.085 | 0.111 | 0.054 | 0.107 | 0.213 | 0.127 | 0.170 | 0.270 | 0.166 | 0.140 | 0.184 | 0.118 | 0.153 | 0.153 | 0.160 |
| | **AM2** | 0.136 | 0.108 | 0.059 | 0.023 | 0.042 | 0.053 | 0.053 | 0.082 | 0.217 | 0.141 | 0.086 | 0.085 | 0.111 | 0.054 | 0.107 | 0.213 | 0.127 | 0.171 | 0.270 | 0.166 | 0.140 | 0.185 | 0.119 | 0.153 | 0.153 | 0.161 |
| | **Comporta** | 0.137 | 0.108 | 0.060 | 0.023 | 0.042 | 0.053 | 0.053 | 0.082 | 0.217 | 0.141 | 0.086 | 0.085 | 0.112 | 0.054 | 0.107 | 0.213 | 0.127 | 0.171 | 0.270 | 0.167 | 0.141 | 0.186 | 0.119 | 0.153 | 0.154 | 0.161 |
| | **TR1** | 0.137 | 0.108 | 0.060 | 0.023 | 0.042 | 0.054 | 0.053 | 0.082 | 0.217 | 0.141 | 0.086 | 0.085 | 0.112 | 0.055 | 0.107 | 0.213 | 0.127 | 0.171 | 0.270 | 0.167 | 0.141 | 0.186 | 0.119 | 0.153 | 0.154 | 0.161 |
| | **TR2** | 0.136 | 0.108 | 0.060 | 0.023 | 0.042 | 0.053 | 0.053 | 0.082 | 0.217 | 0.141 | 0.086 | 0.085 | 0.112 | 0.054 | 0.107 | 0.213 | 0.127 | 0.171 | 0.270 | 0.167 | 0.140 | 0.186 | 0.119 | 0.153 | 0.154 | 0.161 |

| | | KS1 | Boylston | Jap308 | Jap212 | Jap120 | Kasumig2 | Kasumig3 | Kasumig5 | Kosa | GuangDong | GuangXi | Laoshan | ShanDong | Mad23PC | Mad24C | E182 | 128S | E1069 | AM2 | Comporta | TR1 | TR2 |
|---|---|---|---|---|---|---|---|---|---|---|---|---|---|---|---|---|---|---|---|---|---|---|---|
| USA | MO1 | 0.209 | 0.322 | 0.561 | 0.571 | 0.553 | 0.589 | 0.442 | 0.531 | 0.533 | 0.645 | 0.617 | 0.643 | 0.628 | 0.489 | 0.450 | 0.496 | 0.483 | 0.501 | 0.544 | 0.576 | 0.586 | 0.573 |
| | MO2 | 0.189 | 0.318 | 0.569 | 0.580 | 0.559 | 0.616 | 0.431 | 0.591 | 0.532 | 0.673 | 0.644 | 0.673 | 0.656 | 0.453 | 0.403 | 0.462 | 0.444 | 0.468 | 0.521 | 0.560 | 0.572 | 0.556 |
| | NE1 | 0.347 | 0.478 | 0.669 | 0.682 | 0.659 | 0.745 | 0.614 | 0.787 | 0.608 | 0.819 | 0.779 | 0.802 | 0.790 | 0.606 | 0.549 | 0.615 | 0.581 | 0.619 | 0.676 | 0.713 | 0.724 | 0.710 |
| | NE2 | 0.309 | 0.453 | 0.611 | 0.625 | 0.600 | 0.548 | 0.427 | 0.608 | 0.416 | 0.758 | 0.695 | 0.724 | 0.708 | 0.414 | 0.350 | 0.425 | 0.354 | 0.432 | 0.496 | 0.542 | 0.557 | 0.539 |
| | NE5 | 0.315 | 0.461 | 0.722 | 0.734 | 0.712 | 0.735 | 0.557 | 0.761 | 0.574 | 0.818 | 0.768 | 0.793 | 0.779 | 0.572 | 0.506 | 0.583 | 0.543 | 0.589 | 0.652 | 0.694 | 0.706 | 0.690 |
| | NE6 | 0.369 | 0.494 | 0.695 | 0.705 | 0.687 | 0.685 | 0.514 | 0.721 | 0.556 | 0.742 | 0.753 | 0.775 | 0.763 | 0.561 | 0.518 | 0.570 | 0.524 | 0.572 | 0.621 | 0.654 | 0.665 | 0.651 |
| | NE9 | 0.270 | 0.415 | 0.514 | 0.525 | 0.505 | 0.497 | 0.429 | 0.570 | 0.392 | 0.611 | 0.525 | 0.553 | 0.537 | 0.381 | 0.341 | 0.388 | 0.362 | 0.391 | 0.436 | 0.469 | 0.480 | 0.466 |
| | NE10 | 0.366 | 0.488 | 0.704 | 0.713 | 0.696 | 0.720 | 0.621 | 0.737 | 0.618 | 0.809 | 0.737 | 0.759 | 0.747 | 0.609 | 0.569 | 0.616 | 0.595 | 0.619 | 0.662 | 0.693 | 0.702 | 0.690 |
| | NE12 | 0.434 | 0.486 | 0.818 | 0.824 | 0.812 | 0.844 | 0.671 | 0.817 | 0.773 | 0.831 | 0.842 | 0.857 | 0.848 | 0.768 | 0.741 | 0.774 | 0.762 | 0.774 | 0.806 | 0.826 | 0.832 | 0.824 |
| | NE13b | 0.265 | 0.394 | 0.685 | 0.696 | 0.675 | 0.722 | 0.565 | 0.715 | 0.617 | 0.765 | 0.723 | 0.750 | 0.735 | 0.582 | 0.527 | 0.591 | 0.574 | 0.597 | 0.652 | 0.690 | 0.701 | 0.687 |
| | NE14 | 0.276 | 0.440 | 0.607 | 0.617 | 0.599 | 0.629 | 0.542 | 0.653 | 0.516 | 0.688 | 0.672 | 0.696 | 0.682 | 0.511 | 0.471 | 0.518 | 0.493 | 0.521 | 0.566 | 0.598 | 0.609 | 0.595 |
| | NE15 | 0.340 | 0.504 | 0.690 | 0.700 | 0.682 | 0.715 | 0.631 | 0.759 | 0.585 | 0.815 | 0.745 | 0.768 | 0.755 | 0.598 | 0.552 | 0.606 | 0.578 | 0.612 | 0.657 | 0.690 | 0.700 | 0.687 |
| | NE19 | 0.278 | 0.386 | 0.687 | 0.698 | 0.677 | 0.768 | 0.591 | 0.715 | 0.659 | 0.818 | 0.770 | 0.793 | 0.780 | 0.621 | 0.567 | 0.630 | 0.619 | 0.635 | 0.688 | 0.724 | 0.735 | 0.721 |
| | NE22 | 0.260 | 0.420 | 0.566 | 0.579 | 0.555 | 0.609 | 0.446 | 0.639 | 0.482 | 0.707 | 0.671 | 0.701 | 0.684 | 0.435 | 0.378 | 0.446 | 0.392 | 0.451 | 0.512 | 0.555 | 0.568 | 0.551 |
| | NE23 | 0.493 | 0.535 | 0.853 | 0.859 | 0.850 | 0.863 | 0.715 | 0.824 | 0.754 | 0.906 | 0.894 | 0.904 | 0.898 | 0.816 | 0.796 | 0.819 | 0.785 | 0.821 | 0.844 | 0.860 | 0.865 | 0.859 |
| | NE24 | 0.507 | 0.562 | 0.928 | 0.932 | 0.926 | 0.942 | 0.833 | 0.934 | 0.858 | 0.951 | 0.946 | 0.953 | 0.949 | 0.899 | 0.879 | 0.903 | 0.880 | 0.904 | 0.922 | 0.933 | 0.936 | 0.932 |
| | Vi9 | 0.410 | 0.519 | 0.790 | 0.797 | 0.783 | 0.814 | 0.754 | 0.830 | 0.708 | 0.873 | 0.848 | 0.863 | 0.855 | 0.740 | 0.705 | 0.745 | 0.718 | 0.749 | 0.784 | 0.808 | 0.815 | 0.805 |
| | VI10 | 0.225 | 0.356 | 0.687 | 0.698 | 0.678 | 0.717 | 0.568 | 0.709 | 0.600 | 0.740 | 0.754 | 0.778 | 0.764 | 0.595 | 0.542 | 0.605 | 0.590 | 0.610 | 0.664 | 0.701 | 0.712 | 0.697 |
| | MA1 | 0.420 | 0.462 | 0.854 | 0.860 | 0.849 | 0.860 | 0.759 | 0.858 | 0.753 | 0.860 | 0.885 | 0.898 | 0.891 | 0.803 | 0.771 | 0.809 | 0.796 | 0.811 | 0.843 | 0.863 | 0.869 | 0.861 |
| | NC1 | 0.180 | 0.338 | 0.638 | 0.651 | 0.626 | 0.698 | 0.531 | 0.685 | 0.595 | 0.740 | 0.739 | 0.767 | 0.751 | 0.547 | 0.480 | 0.559 | 0.540 | 0.565 | 0.632 | 0.676 | 0.688 | 0.671 |
| | NC2 | 0.158 | 0.305 | 0.550 | 0.563 | 0.541 | 0.624 | 0.508 | 0.599 | 0.549 | 0.685 | 0.665 | 0.692 | 0.676 | 0.495 | 0.445 | 0.503 | 0.487 | 0.509 | 0.561 | 0.600 | 0.611 | 0.596 |
| | NY1 | 0.186 | 0.350 | 0.716 | 0.729 | 0.705 | 0.781 | 0.664 | 0.780 | 0.678 | 0.831 | 0.802 | 0.826 | 0.812 | 0.630 | 0.555 | 0.642 | 0.631 | 0.648 | 0.714 | 0.755 | 0.767 | 0.752 |
| | GA1 | 0.182 | 0.389 | 0.798 | 0.810 | 0.790 | 0.852 | 0.685 | 0.843 | 0.705 | 0.896 | 0.865 | 0.883 | 0.873 | 0.702 | 0.625 | 0.714 | 0.683 | 0.721 | 0.781 | 0.817 | 0.826 | 0.814 |
| | GA2 | 0.215 | 0.340 | 0.581 | 0.593 | 0.571 | 0.662 | 0.492 | 0.647 | 0.589 | 0.715 | 0.682 | 0.709 | 0.693 | 0.522 | 0.472 | 0.531 | 0.520 | 0.536 | 0.589 | 0.628 | 0.639 | 0.624 |
| | GA3 | 0.127 | 0.295 | 0.725 | 0.739 | 0.713 | 0.798 | 0.614 | 0.777 | 0.676 | 0.849 | 0.824 | 0.847 | 0.834 | 0.652 | 0.567 | 0.666 | 0.643 | 0.674 | 0.742 | 0.783 | 0.794 | 0.780 |
| | GA4 | 0.154 | 0.285 | 0.600 | 0.613 | 0.588 | 0.669 | 0.549 | 0.659 | 0.577 | 0.744 | 0.663 | 0.694 | 0.676 | 0.525 | 0.464 | 0.536 | 0.521 | 0.542 | 0.604 | 0.647 | 0.659 | 0.642 |
| | KS1 | | 0.246 | 0.619 | 0.633 | 0.609 | 0.672 | 0.558 | 0.658 | 0.590 | 0.737 | 0.691 | 0.720 | 0.703 | 0.525 | 0.466 | 0.535 | 0.529 | 0.542 | 0.600 | 0.642 | 0.654 | 0.638 |
| | Boylston | 0.157 | | 0.682 | 0.690 | 0.675 | 0.701 | 0.610 | 0.663 | 0.646 | 0.741 | 0.716 | 0.737 | 0.725 | 0.625 | 0.592 | 0.631 | 0.631 | 0.635 | 0.671 | 0.698 | 0.706 | 0.696 |
| Japan | Jap308 | 0.218 | 0.357 | | 0 | 0 | 0.992 | 0.883 | 0.995 | 0.821 | 0.997 | 0.992 | 0.993 | 0.993 | 0.017 | 0.017 | 0.017 | 0.020 | 0.017 | 0.017 | 0.017 | 0.017 | 0.017 |
| | Jap212 | 0.218 | 0.357 | 0 | | 0 | 0.993 | 0.886 | 0.995 | 0.827 | 0.997 | 0.992 | 0.993 | 0.993 | 0.017 | 0.017 | 0.017 | 0.020 | 0.017 | 0.017 | 0.017 | 0.017 | 0.017 |
| | Jap120 | 0.218 | 0.357 | 0 | 0 | | 0.992 | 0.879 | 0.995 | 0.817 | 0.997 | 0.992 | 0.992 | 0.992 | 0.017 | 0.017 | 0.017 | 0.020 | 0.017 | 0.017 | 0.017 | 0.017 | 0.017 |
| | Kasumig2 | 0.216 | 0.302 | 0.017 | 0.017 | 0.017 | | 0.864 | 0.985 | 0.627 | 0.996 | 0.992 | 0.992 | 0.992 | 0.017 | 0.017 | 0.017 | 0.017 | 0.017 | 0.017 | 0.017 | 0.017 | 0.017 |
| | Kasumig3 | 0.205 | 0.271 | 0.094 | 0.094 | 0.094 | 0.054 | | 0.835 | 0.746 | 0.866 | 0.895 | 0.905 | 0.899 | 0.031 | 0.031 | 0.031 | 0.031 | 0.031 | 0.031 | 0.031 | 0.031 | 0.031 |
| | Kasumig5 | 0.197 | 0.236 | 0.038 | 0.038 | 0.038 | 0.004 | 0.033 | | 0.789 | 0.996 | 0.995 | 0.995 | 0.995 | 0.038 | 0.038 | 0.038 | 0.038 | 0.038 | 0.038 | 0.038 | 0.038 | 0.038 |
| | Kosa | 0.215 | 0.306 | 0.036 | 0.036 | 0.036 | 0.004 | 0.058 | 0.017 | | 0.875 | 0.828 | 0.841 | 0.834 | 0.018 | 0.018 | 0.018 | 0.018 | 0.018 | 0.018 | 0.018 | 0.018 | 0.018 |
| China | GuangDong | 0.297 | 0.366 | 0.106 | 0.106 | 0.106 | 0.068 | 0.040 | 0.068 | 0.041 | | 0.997 | 0.997 | 0.997 | 0.068 | 0.068 | 0.068 | 0.068 | 0.068 | 0.068 | 0.068 | 0.068 | 0.068 |
| | GuangXi | 0.265 | 0.373 | 0.017 | 0.017 | 0.017 | 0.017 | 0.094 | 0.038 | 0.034 | 0.106 | | 0 | 0 | 0.017 | 0.017 | 0.017 | 0.020 | 0.017 | 0.017 | 0.017 | 0.017 | 0.017 |
| | Laoshan | 0.265 | 0.373 | 0.017 | 0.017 | 0.017 | 0.017 | 0.094 | 0.038 | 0.034 | 0.106 | 0 | | 0 | 0.017 | 0.017 | 0.017 | 0.020 | 0.017 | 0.017 | 0.017 | 0.017 | 0.017 |
| | ShanDong | 0.265 | 0.373 | 0.017 | 0.017 | 0.017 | 0.017 | 0.094 | 0.038 | 0.034 | 0.106 | 0 | 0 | | 0.017 | 0.017 | 0.017 | 0.020 | 0.017 | 0.017 | 0.017 | 0.017 | 0.017 |
| Portugal | Mad23PC | 0.216 | 0.366 | 0.992 | 0.992 | 0.992 | 0.992 | 0.744 | 0.995 | 0.726 | 0.996 | 0.992 | 0.992 | 0.992 | | 0 | 0 | 0.295 | 0 | 0 | 0 | 0 | 0 |
| | Mad24C | 0.214 | 0.366 | 0.992 | 0.992 | 0.992 | 0.992 | 0.718 | 0.995 | 0.701 | 0.996 | 0.992 | 0.992 | 0.992 | 0 | | 0 | 0.243 | 0 | 0 | 0 | 0 | 0 |
| | E182 | 0.217 | 0.367 | 0.992 | 0.992 | 0.992 | 0.992 | 0.750 | 0.995 | 0.730 | 0.996 | 0.992 | 0.992 | 0.992 | 0 | 0 | | 0.304 | 0 | 0 | 0 | 0 | 0 |
| | 128S | 0.216 | 0.367 | 0.915 | 0.919 | 0.912 | 0.913 | 0.687 | 0.942 | 0.667 | 0.962 | 0.926 | 0.934 | 0.930 | 0.001 | 0.001 | 0.001 | | 0.312 | 0.362 | 0.398 | 0.414 | 0.398 |
| | E1069 | 0.217 | 0.367 | 0.992 | 0.992 | 0.992 | 0.992 | 0.752 | 0.995 | 0.733 | 0.996 | 0.992 | 0.992 | 0.992 | 0 | 0 | 0 | 0.001 | | 0 | 0 | 0 | 0 |
| | AM2 | 0.218 | 0.367 | 0.992 | 0.992 | 0.992 | 0.992 | 0.782 | 0.995 | 0.760 | 0.996 | 0.992 | 0.992 | 0.992 | 0 | 0 | 0 | 0.001 | 0 | | 0 | 0 | 0 |
| | Comporta | 0.218 | 0.367 | 0.992 | 0.992 | 0.992 | 0.992 | 0.801 | 0.995 | 0.779 | 0.996 | 0.992 | 0.992 | 0.992 | 0 | 0 | 0 | 0.001 | 0 | 0 | | 0 | 0 |
| | TR1 | 0.218 | 0.367 | 0.992 | 0.992 | 0.992 | 0.992 | 0.809 | 0.995 | 0.786 | 0.996 | 0.992 | 0.992 | 0.992 | 0 | 0 | 0 | 0.001 | 0 | 0 | 0 | | 0 |
| | TR2 | 0.218 | 0.367 | 0.992 | 0.992 | 0.992 | 0.992 | 0.800 | 0.995 | 0.778 | 0.996 | 0.992 | 0.992 | 0.992 | 0 | 0 | 0 | 0.001 | 0 | 0 | 0 | 0 | |

**Table S2:** Results of all ABC Random Forest analyses obtained with DIYABC Random Forest v1.0 (Collin et al. 2021). Number of votes for each scenario are based on 1,000 decision trees per analysis. The posterior probabilities and number of votes of the selected scenario in each analysis are shown in bold.

| Analyses - Scenarios | Prior error rate | | | | Random forest votes (posterior probability) | | | |
|---|---|---|---|---|---|---|---|---|
| | Main 30G/y | Alt. 30G/y | Main 15G/y | Main 45G/y | Main 30G/y | Alt. 30G/y | Main 15G/y | Main 45G/y |
| *Step 1 – Japan (301 SuSts)* | 0.273 | 0.253 | 0.250 | 0.302 | | | | |
| S1: Nat→Japan1 ; Nat→Japan2 ; Nat→Japan3 | | | | | 338 | 321 | **423 (0.774)** | 266 |
| S2: Nat→Japan1→Japan2→Japan3 | | | | | 14 | 7 | 11 | 10 |
| S3: Nat→Japan1→Japan2 ; Nat→Japan3 | | | | | 112 | 100 | 102 | 72 |
| S4: Nat→Japan1→Japan3 ; Nat→Japan2 | | | | | **452 (0.711)** | **501 (0.672)** | 406 | **528 (0.651)** |
| S5: Nat→Japan1 ; Japan1→Japan2 ; Japan1→Japan3 | | | | | 71 | 57 | 41 | 101 |
| S6: Nat→Japan1 ; Nat→Japan2→Japan3 | | | | | 6 | 11 | 7 | 15 |
| S7: Nat→Japan1→Japan3→Japan2 | | | | | 2 | 1 | 3 | 7 |
| S8: Nat→Japan1 ; Nat→Japan3→Japan2 | | | | | 5 | 2 | 7 | 1 |
| *Step 2 – China (204 SuSts)* | 0.171 | 0.164 | 0.160 | 0.181 | | | | |
| S1: Native→China1 ; Native→China2 | | | | | **960 (0.963)** | **960 (0.957)** | **986 (0.984)** | **929 (0.932)** |
| S2: Native→China1→China2 | | | | | 30 | 26 | 10 | 65 |
| S3: Native→China2→China1 | | | | | 10 | 14 | 4 | 6 |
| *Step 3 – Japan/China (576 SuSts)* | 0.338 | 0.302 | 0.354 | 0.357 | | | | |
| S1: Nat → Japan1 →Japan3 ; Nat → Japan2 ; Nat → China1 ; Nat → China2 | | | | | 44 | 61 | 89 | 34 |
| S2: Nat → Japan1 →Japan3 ; Nat → Japan2 ; Japan1 → China1 ; Nat → China2 | | | | | **246 (0.573)** | **272 (0.586)** | **206 (0.575)** | 157 |
| S3: Nat → Japan1 →Japan3 ; Nat → Japan2 ; Nat → China1 ; Japan1 → China2 | | | | | 18 | 16 | 16 | 10 |
| S4: Nat → Japan1 →Japan3 ; Nat → Japan2 ; Japan2 → China1 ; Nat → China2 | | | | | 11 | 8 | 8 | 9 |
| S5: Nat → Japan1 →Japan3 ; Nat → Japan2 ; Nat → China1 ; Japan2 → China2 | | | | | 29 | 41 | 33 | 41 |
| S6: Nat → Japan1 →Japan3 ; Nat → Japan2 ; Japan3 → China1 ; Nat → China2 | | | | | 190 | 194 | 163 | **213 (0.551)** |
| S7: Nat → Japan1 →Japan3 ; Nat → Japan2 ; Nat → China1 ; Japan3 → China2 | | | | | 4 | 4 | 6 | 2 |
| S8: Nat → Japan1 →Japan3 ; Nat → Japan2 ; Japan1 → China1 ; Japan1 → China2 | | | | | 25 | 35 | 35 | 41 |
| S9: Nat → Japan1 →Japan3 ; Nat → Japan2 ; Japan2 → China1 ; Japan2 → China2 | | | | | 3 | 3 | 4 | 4 |
| S10: Nat → Japan1 →Japan3 ; Nat → Japan2 ; Japan3 → China1 ; Japan3 → China2 | | | | | 6 | 7 | 5 | 11 |
| S11 : Nat → Japan1 →Japan3 ; Nat → Japan2 ; Japan1 → China1 ; Japan2 → China2 | | | | | 158 | 134 | 97 | 172 |
| S12 : Nat → Japan1 →Japan3 ; Nat → Japan2 ; Japan1 → China1 ; Japan3 → China2 | | | | | 12 | 5 | 10 | 17 |
| S13 : Nat → Japan1 →Japan3 ; Nat → Japan2 ; Japan2 → China1 ; Japan1 → China2 | | | | | 1 | 4 | 1 | 2 |
| S14 : Nat → Japan1 →Japan3 ; Nat → Japan2 ; Japan2 → China1 ; Japan3 → China2 | | | | | 0 | 0 | 3 | 2 |
| S15 : Nat → Japan1 →Japan3 ; Nat → Japan2 ; Japan3 → China1 ; Japan1 → China2 | | | | | 34 | 33 | 49 | 49 |
| S16 : Nat → Japan1 →Japan3 ; Nat → Japan2 ; Japan3 → China1 ; Japan2 → China2 | | | | | 143 | 94 | 70 | 184 |
| S17: Nat → Japan1 ; Nat →Japan2 ; Nat → China1 → Japan3 ; Nat → China2 | | | | | 64 | 77 | 192 | 49 |
| S18: Nat → Japan1 ; Nat →Japan2 ; Nat → China1 ; Nat → China2 → Japan3 | | | | | 1 | 1 | 3 | 0 |
| S19: Nat → Japan1 → Japan3 ; Nat → China1 → Japan2 ; Nat → China2 | | | | | 1 | 1 | 1 | 2 |
| S20: Nat → Japan1 → Japan3 ; Nat → China1 ; Nat → China2 → Japan2 | | | | | 0 | 0 | 0 | 0 |
| S21: Nat → Japan1 ; Nat → China1 → Japan2 ; China1 → Japan3 ; Nat → China2 | | | | | 8 | 10 | 8 | 0 |
| S22: Nat → Japan1 ; Nat → China1 ; Nat → China2 → Japan2 ; China2 → Japan3 | | | | | 0 | 0 | 0 | 0 |

| | | | | | | | | |
|---|---|---|---|---|---|---|---|---|
| S23: Nat → Japan1 ; Nat → China1 → Japan2 ; Nat → China2 → Japan3 | | | | | 0 | 0 | 0 | 0 |
| S24: Nat → Japan1 ; Nat → China1 → Japan3 ; Nat → China2 → Japan2 | | | | | 2 | 0 | 1 | 1 |
| *Step 4 – Japan/China admixture hypothesis (576 SuSts)* | 0.311 | 0.306 | 0.290 | 0.326 | | | | |
| S1: Scenario 2 of step 3 | | | | | **429 (0.567)** | **453 (0.531)** | **385 (0.506)** | **409 (0.564)** |
| S2: Nat → Japan1 →Japan3 ; Japan1 → China1 ; Nat → China2 ; Nat+China2 → Japan2 | | | | | 390 | 334 | 370 | 388 |
| S3: Nat → Japan1 →Japan3 ; Japan1 → China1 ; Nat → China2 ; Japan1+China2 → Japan2 | | | | | 181 | 213 | 245 | 203 |
| *Step 5 – Portugal (760 SuSts)* | 0.208 | 0.182 | 0.249 | 0.200 | | | | |
| S1: Nat → Japan1 →Japan3 ; Nat → Japan2 ; Japan1 → China1 ; Nat → China2 ; Nat → Port | | | | | 218 | 231 | 241 | 206 |
| S2: Nat → Japan1 →Japan3 ; Nat → Japan2 ; Japan1 → China1 ; Nat → China2 ; Japan1 → Port | | | | | **329 (0.682)** | **323 (0.675)** | **269 (0.663)** | **284 (0.649)** |
| S3: Nat → Japan1 →Japan3 ; Nat → Japan2 → Port ; Japan1 → China1 ; Nat → China2 | | | | | 96 | 101 | 95 | 108 |
| S4: Nat → Japan1 →Japan3 → Port ; Nat → Japan2 ; Japan1 → China1 ; Nat → China2 | | | | | 154 | 161 | 194 | 182 |
| S5: Nat → Japan1 →Japan3 ; Nat → Japan2 ; Japan1 → China1 → Port ; Nat → China2 | | | | | 170 | 148 | 158 | 177 |
| S6: Nat → Japan1 →Japan3 ; Nat → Japan2 ; Japan1 → China1 ; Nat → China2 → Port | | | | | 21 | 28 | 29 | 35 |
| S7: Nat → Japan1 →Japan3 ; Nat → Japan2 ; Native → Port → China1 ; Nat → China2 | | | | | 2 | 1 | 8 | 1 |
| S8: Nat → Japan1 →Japan3 ; Nat → Japan2 ; Japan1 → China1 ; Nat → Port → China2 | | | | | 10 | 7 | 6 | 7 |
| *Step 6 – Invasive bridgehead hypothesis (760 SuSts)* | 0.356 | 0.343 | 0.327 | 0.368 | | | | |
| S1: Scenario 2 of step 5 | | | | | **428 (0.567)** | **432 (0.578)** | **436 (0.532)** | **405 (0.566)** |
| S2: As S1, but Japan1, Japan2 and China2 all derive from an invasive ghost population | | | | | 190 | 216 | 148 | 274 |
| S3: As S1, but Japan1, Japan2 and China2 all derive from a single native ghost population | | | | | 382 | 352 | 416 | 321 |

**Table S3:** ABC Model checking results for the final selected scenario (Table S2, step 6; Figure 3A), and for each prior set and time scale. For each analysis, the total number of summary statistics is 760. FDR stand for "false discovery rate".

| Prior set | Nb of generation/year | Nb of SuSts with P < 0.05 or P > 0.95 | Prop of SuSts with P < 0.05 or P > 0.95 | Nb after FDR correction | Prop after FDR correction |
|---|---|---|---|---|---|
| Main | 30 | 56 | 0.074 | 0 | 0.000 |
| Alternate | 30 | 52 | 0.068 | 0 | 0.000 |
| Main | 15 | 41 | 0.054 | 0 | 0.000 |
| Main | 45 | 19 | 0.025 | 0 | 0.000 |

**Figure S1:** Worldwide location of the sampling sites for the *Bursaphelenchus xylophilus* specimens used in this study. Four different geographic areas have been investigated: the USA, as part of the native area (A), and Japan, mainland Protugal and Madeira Island, and China as invaded areas (B, C and D, respectively). For details on the samples, see Table 1.

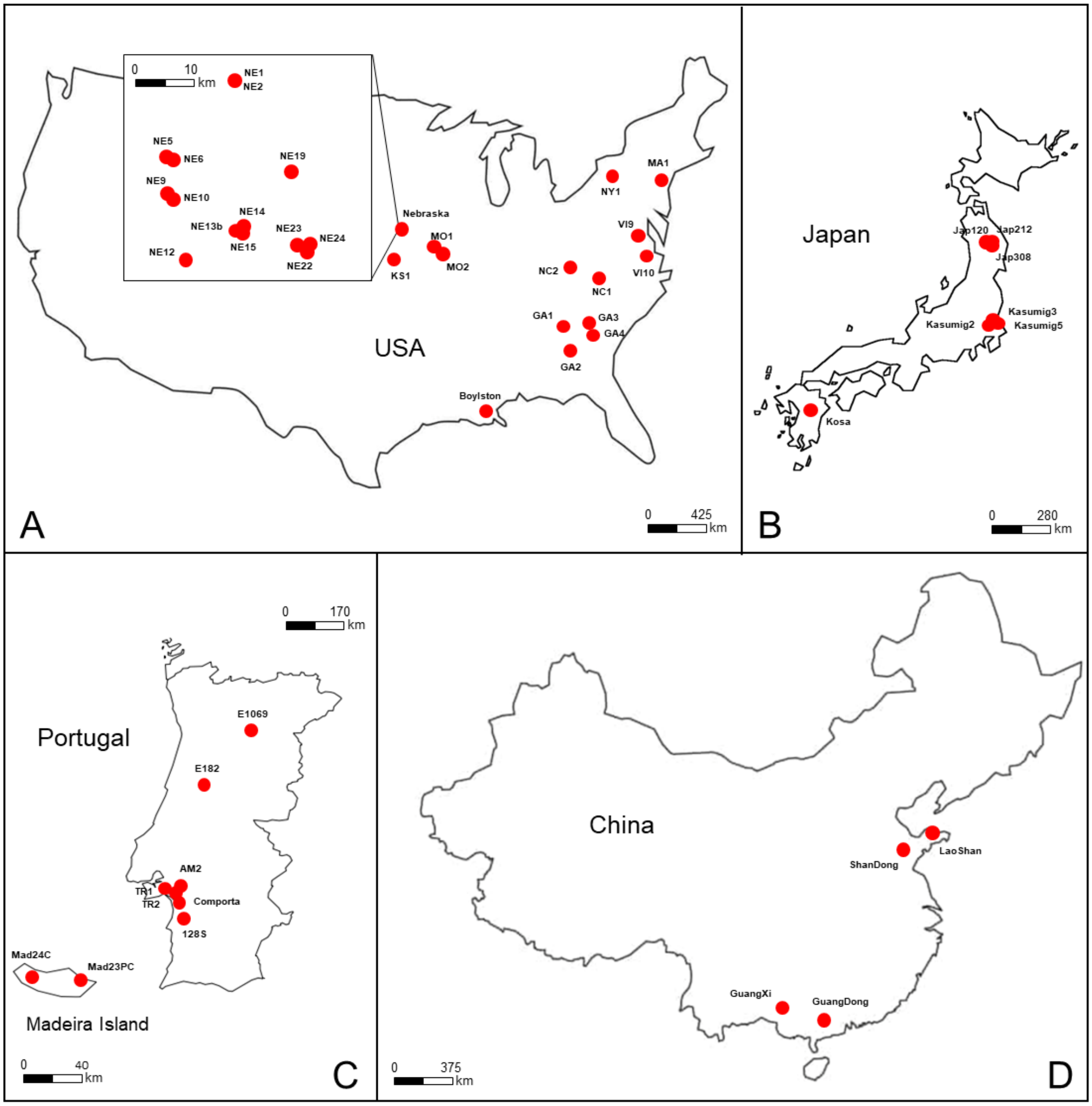

**Figure S2:** illustration of a few competing evolutionary scenarios simulated for the ABC-RF analyses. For each scenario, a graphical representation and the specific DIYABC Random Forest v1.0 code (with associated conditions) are provided. Details about parameters and prior distributions can be found in Table 2.

### (A) Step 1, Scenario 4

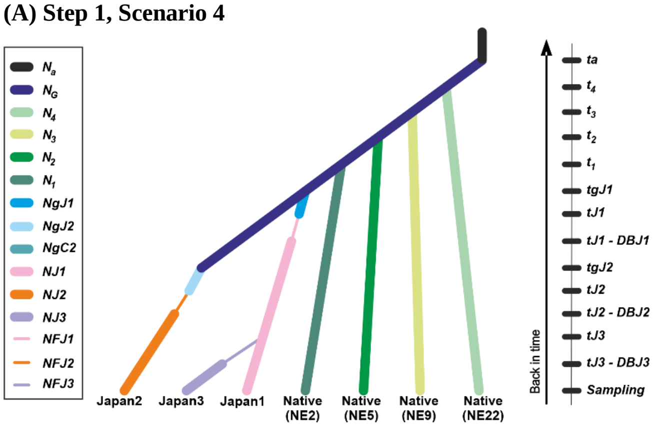


**DIYABC Random Forest v1.0 code:**

```
N1 NJ1 NJ2 NJ3 N2 N3 N4 NG
0 sample 1
0 sample 2
0 sample 3
0 sample 4
0 sample 5
0 sample 6
0 sample 7
tJ3-DBJ3 varNe 4 NF3
tJ3 merge 2 4
tJ2-DBJ2 varNe 3 NFJ2
tJ2 VarNe 3 NgJ2
tgJ2 merge 8 3
tJ1-DBJ1 varNe 2 NFJ1
tJ1 VarNe 2 NgJ1
tgJ1 merge 8 2
t1 merge 8 1
t2 merge 8 5
t3 merge 8 6
t4 merge 8 7
ta VarNe 8 Na
```

**Conditions:**

```
Conditions:
tgJ1>tJ1
tgJ2>tJ2
ta>tgJ1
ta>tgJ2
ta>t1
ta>t2
ta>t3
ta>t4
```

### (B) Step 3, Scenario 2

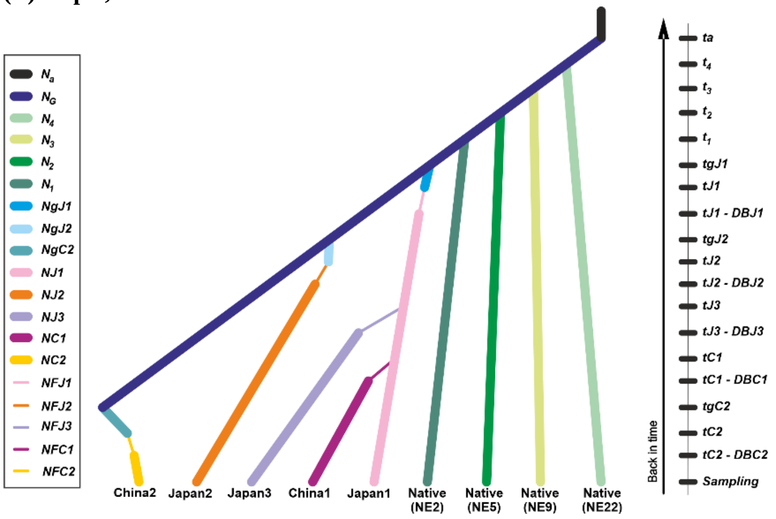


```
N1 NJ1 NJ2 NJ3 NC1 NC2 N2 N3 N4 NG
0 sample 1
0 sample 2
0 sample 3
0 sample 4
0 sample 5
0 sample 6
0 sample 7
0 sample 8
0 sample 9
tC2-DBC2 varNe 6 NFC2
tC2 VarNe 6 NgC2
tgC2 merge 10 6
tC1-DBC1 varNe 5 NFC1
tC1 merge 2 5
tJ3-DBJ3 varNe 4 NF3
tJ3 merge 2 4
tJ2-DBJ2 varNe 3 NFJ2
tJ2 VarNe 3 NgJ2
tgJ2 merge 8 3
tJ1-DBJ1 varNe 2 NFJ1
tJ1 VarNe 2 NgJ1
tgJ1 merge 8 2
t1 merge 8 1
t2 merge 8 5
t3 merge 8 6
t4 merge 8 7
ta VarNe 8 Na
```

```
tgJ1>tJ1
tgJ2>tJ2
tgC2>tC2
ta>tgJ1
ta>tgJ2
ta>tgC2
ta>t1
ta>t2
ta>t3
ta>t4
```

### (C) Step 6, Scenario 1

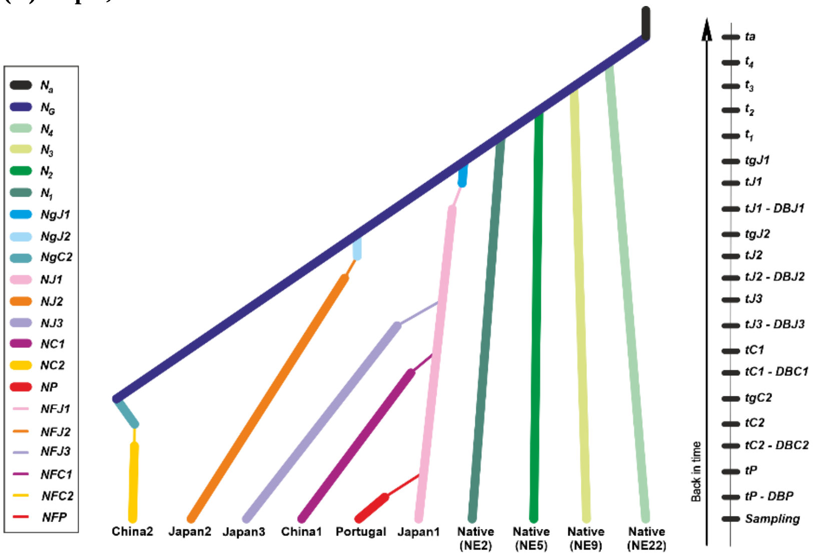


```
N1 NJ1 NJ2 NJ3 NC1 NC2 NP N2 N3 N4 NG
0 sample 1
0 sample 2
0 sample 3
0 sample 4
0 sample 5
0 sample 6
0 sample 7
0 sample 8
0 sample 9
0 sample 10
tP-DBP VarNe 7 NFP
tP merge 2 7
tC2-DBC2 varNe 6 NFC2
tC2 VarNe 6 NgC2
tgC2 merge 10 6
tC1-DBC1 varNe 5 NFC1
tC1 merge 2 5
tJ3-DBJ3 varNe 4 NF3
tJ3 merge 2 4
tJ2-DBJ2 varNe 3 NFJ2
tJ2 VarNe 3 NgJ2
tgJ2 merge 8 3
tJ1-DBJ1 varNe 2 NFJ1
tJ1 VarNe 2 NgJ1
tgJ1 merge 8 2
t1 merge 8 1
t2 merge 8 5
t3 merge 8 6
t4 merge 8 7
ta VarNe 8 Na
```

```
tgJ1>tJ1
tgJ2>tJ2
tgC2>tC2
ta>tgJ1
ta>tgJ2
ta>tgC2
ta>t1
ta>t2
ta>t3
ta>t4
```

**Figure S3: Genetic structure of the American samples of PWN.** (A) Mean log-likelihood and Δ$K$ (Evanno et al., 2005, *Molecular Ecology*, 14:2611-2620) for each value of $K$ tested. (B) Bar plots of the coefficients of co-ancestry obtained in the Structure analysis with several values of $K$. Each bar corresponds to one individual nematode and each cluster is represented with a particular color. The two most frequent clustering pattern are shown, when the major one represents less than 15 runs out of the 20 performed.

(A)

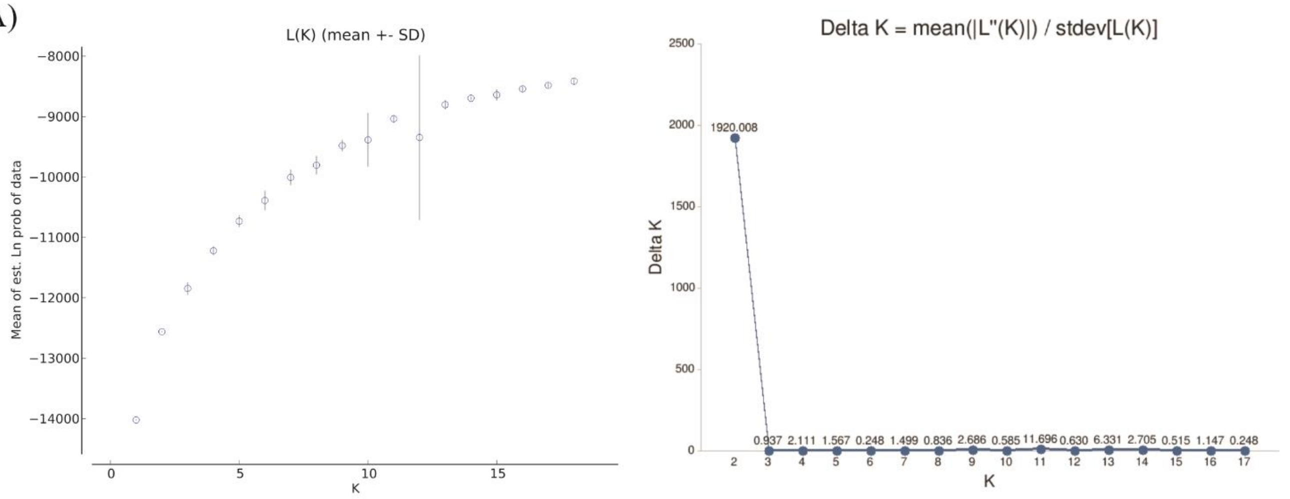


(B)

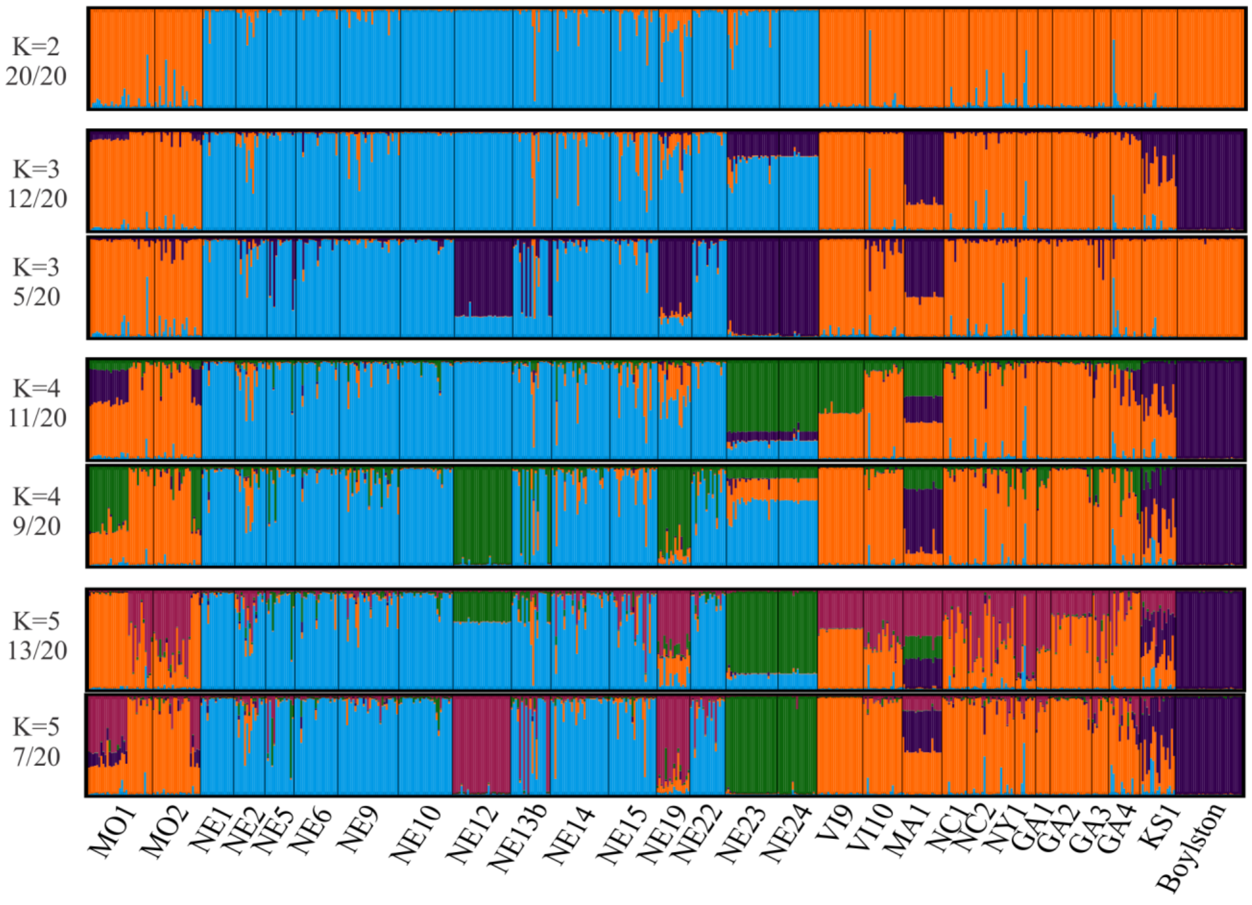

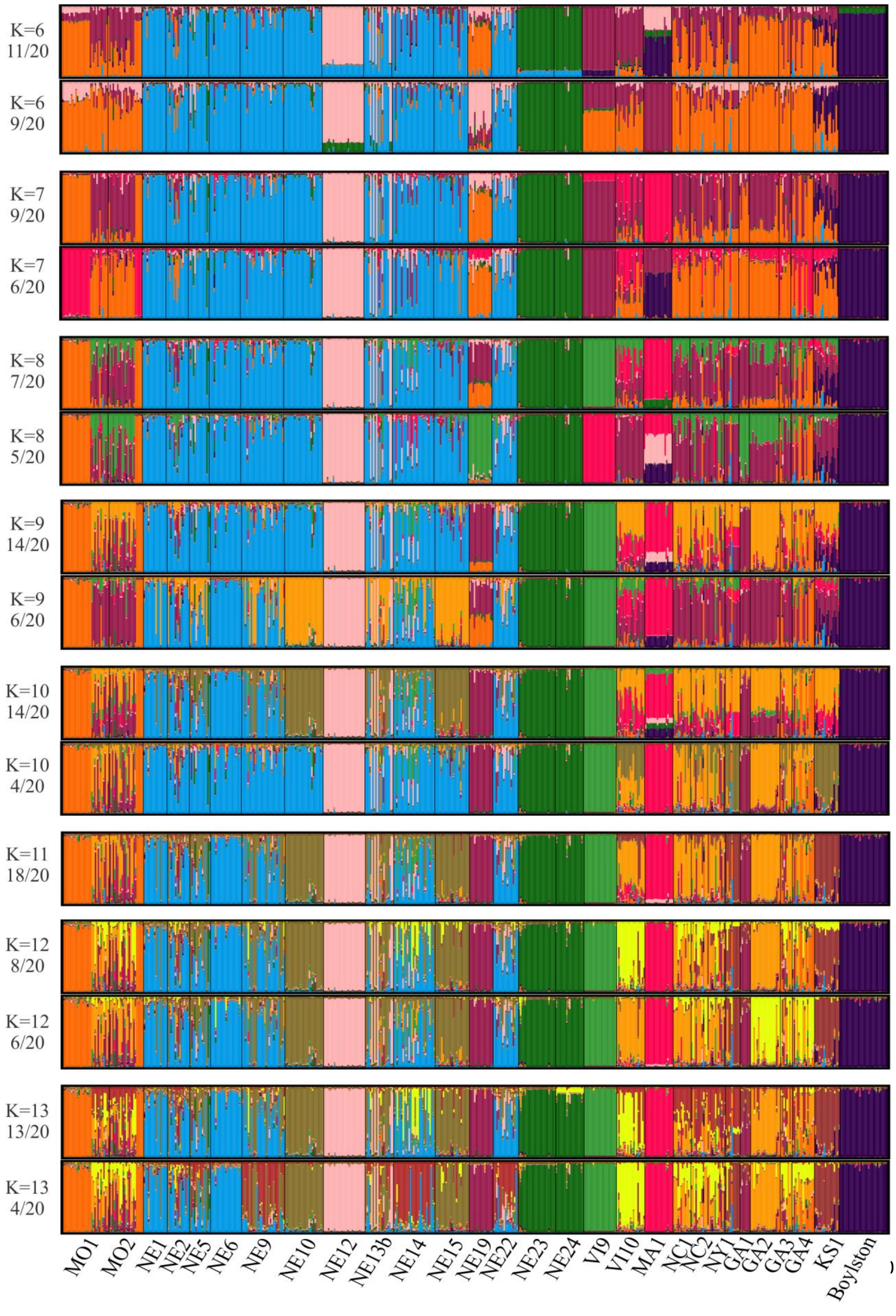
K=6
11/20
K=6
9/20
K=7
9/20
K=7
6/20
K=8
7/20
K=8
5/20
K=9
14/20
K=9
6/20
K=10
14/20
K=10
4/20
K=11
18/20
K=12
8/20
K=12
6/20
K=13
13/20
K=13
4/20
MO1
MO2
NE1
NE2
NE5
NE6
NE9
NE10
NE12
NE13b
NE14
NE15
NE19
NE22
NE23
NE24
VI9
VI10
MA1
NC1
NC2
NY1
GA1
GA2
GA3
GA4
KS1
Boylston

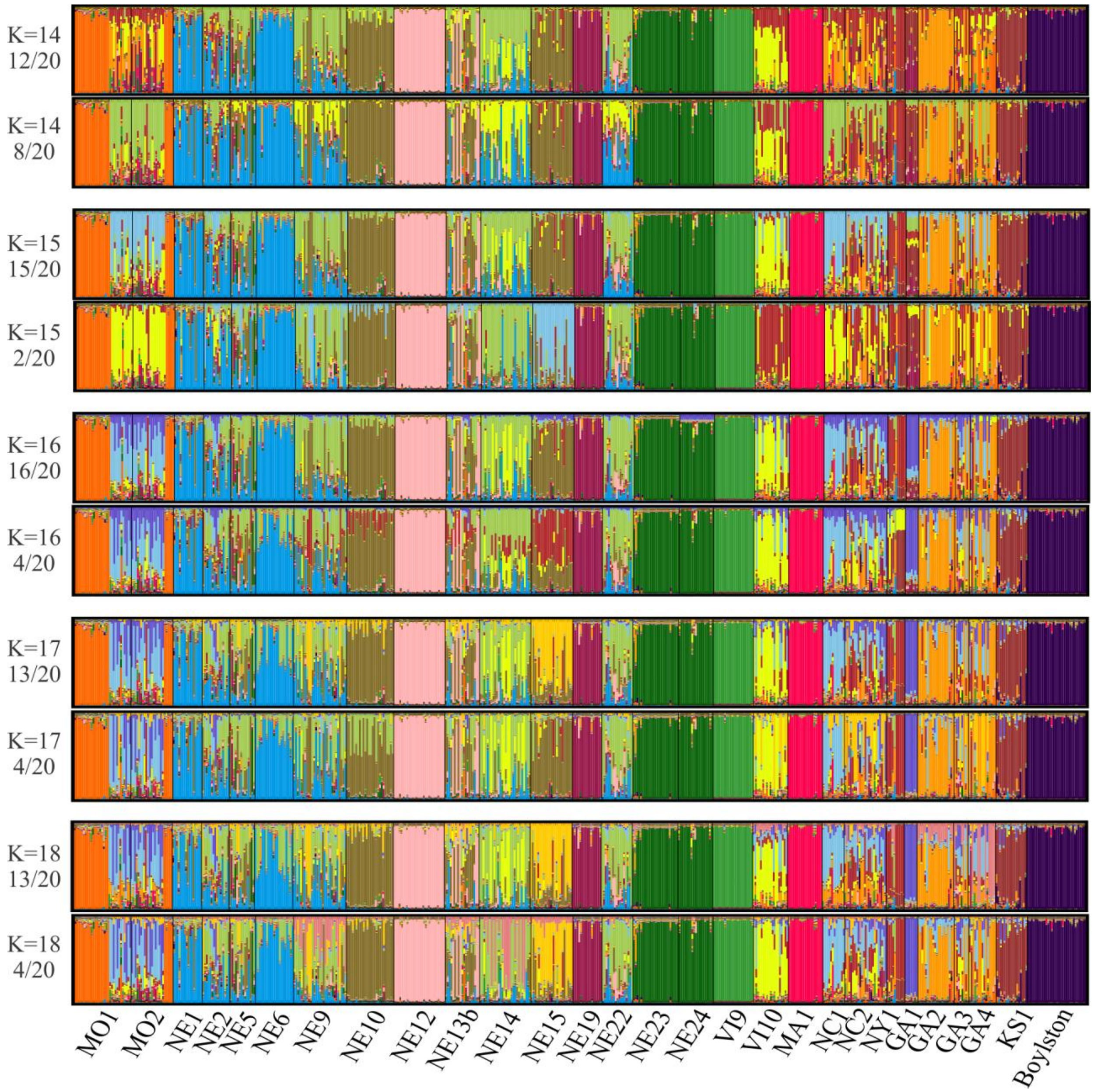
K=14
12/20
K=14
8/20
K=15
15/20
K=15
2/20
K=16
16/20
K=16
4/20
K=17
13/20
K=17
4/20
K=18
13/20
K=18
4/20
MO1
MO2
NE1
NE2
NE5
NE6
NE9
NE10
NE12
NE13b
NE14
NE15
NE19
NE22
NE23
NE24
VI9
VI10
MA1
NC1
NC2
NY1
GA1
GA2
GA3
GA4
KS1
Boylston

**Figure S4: Genetic structure of the Chinese samples of PWN.** Bar plots of the coefficients of co-ancestry obtained in the Structure analysis with several values of K. Each bar corresponds to one individual nematode and each cluster is represented with a particular color. Only one clustering pattern per value of K was identified by Clump.

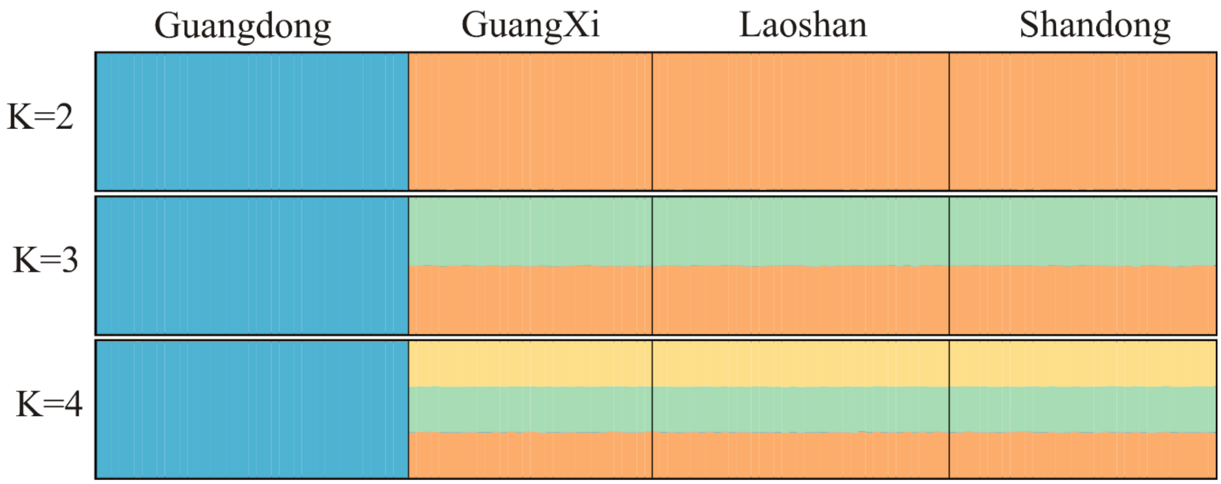

**Figure S5: (A) Genetic structure of the Japanese samples of PWN. (B) Genetic structure of the European samples of PWN.** Bar plots of the coefficients of co-ancestry obtained in the Structure analysis with several values of K. Each bar corresponds to one individual nematode and each cluster is represented with a particular color. Only one clustering pattern per value of K was identified by Clump. Plots taken from Mallez et al (2015, *Biological Invasions*, 17:1199–1213).

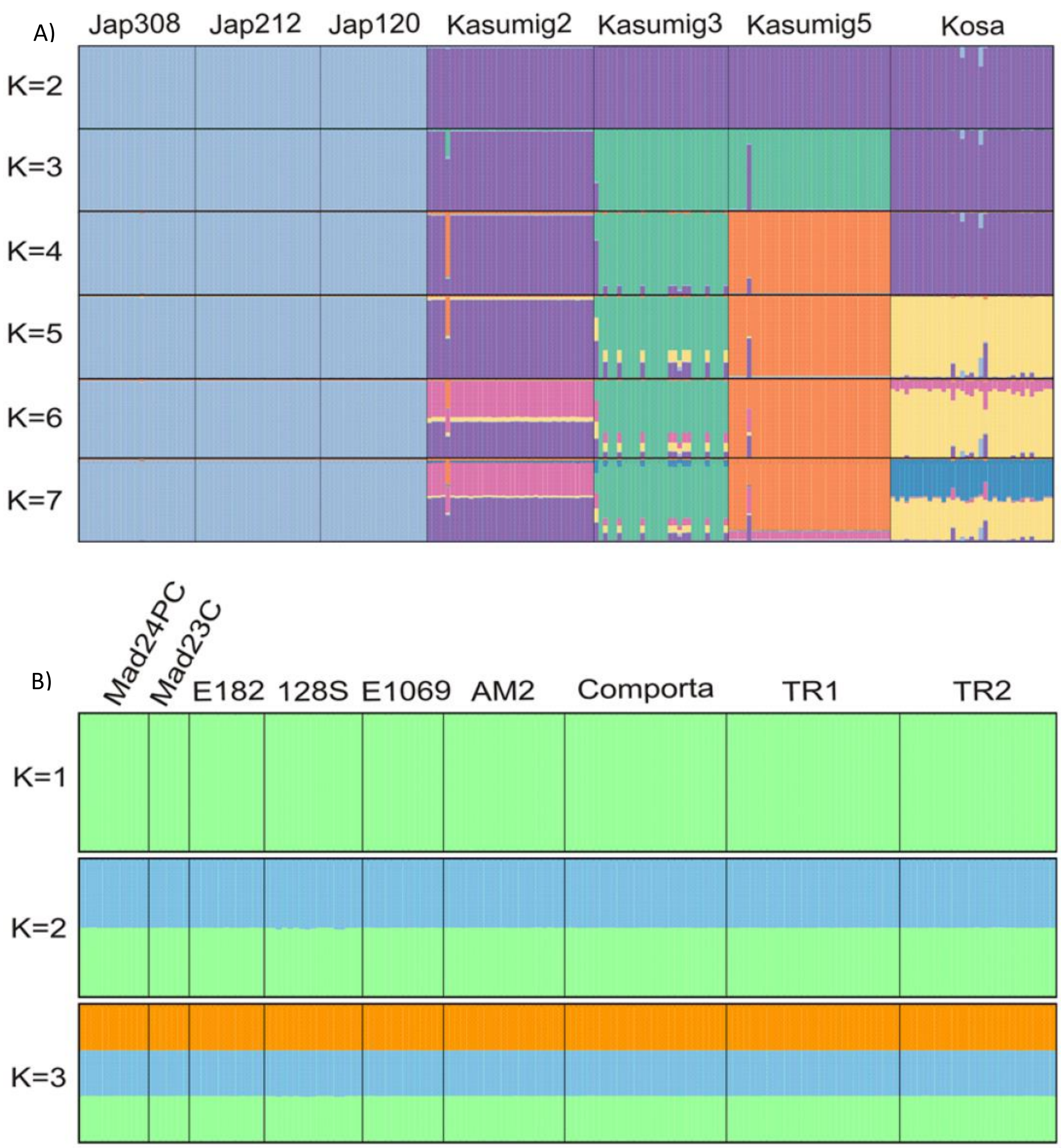

**Figure S6: Genetic structure of the PWN samples from all areas.** (A) Mean log-likelihood and ΔK (Evanno et al., 2005, *Molecular Ecology*, 14:2611-2620) for each value of K tested. (B) The second most frequent clustering patterns is shown when the most frequent (the major, shown in Fig. 3) one represents less than 15 runs out of the 20 performed.

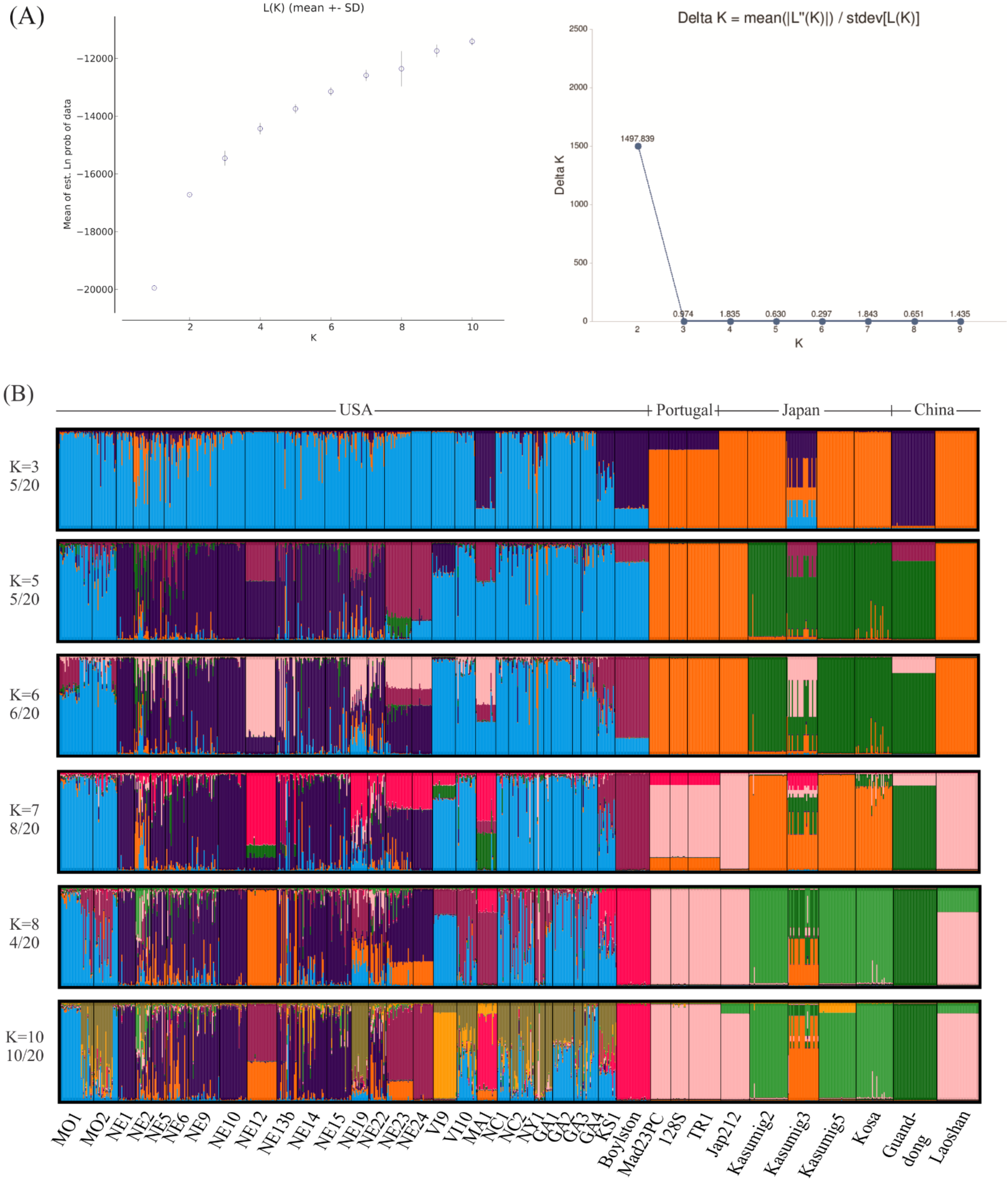

**Figure S7: Mean individual assignment likelihoods of the invasive samples (Japanese, Chinese, and Portuguese) to all their potential sources.** Top panel - Japanese invasive samples considered, implying that only one potential source exits: the native area, USA. Bottom panel – Chinese and Portuguese invasive samples considered, implying that two (USA, Japan) and three (USA, Japan and China) potential sources exist, respectively. Only one sample among the “undifferentiated” ones was kept per invaded area, in this analysis for simplification: Jap212 for Japan3, Laoshan for China1, TR1 for Portugal and Mad23PC for Madeira.

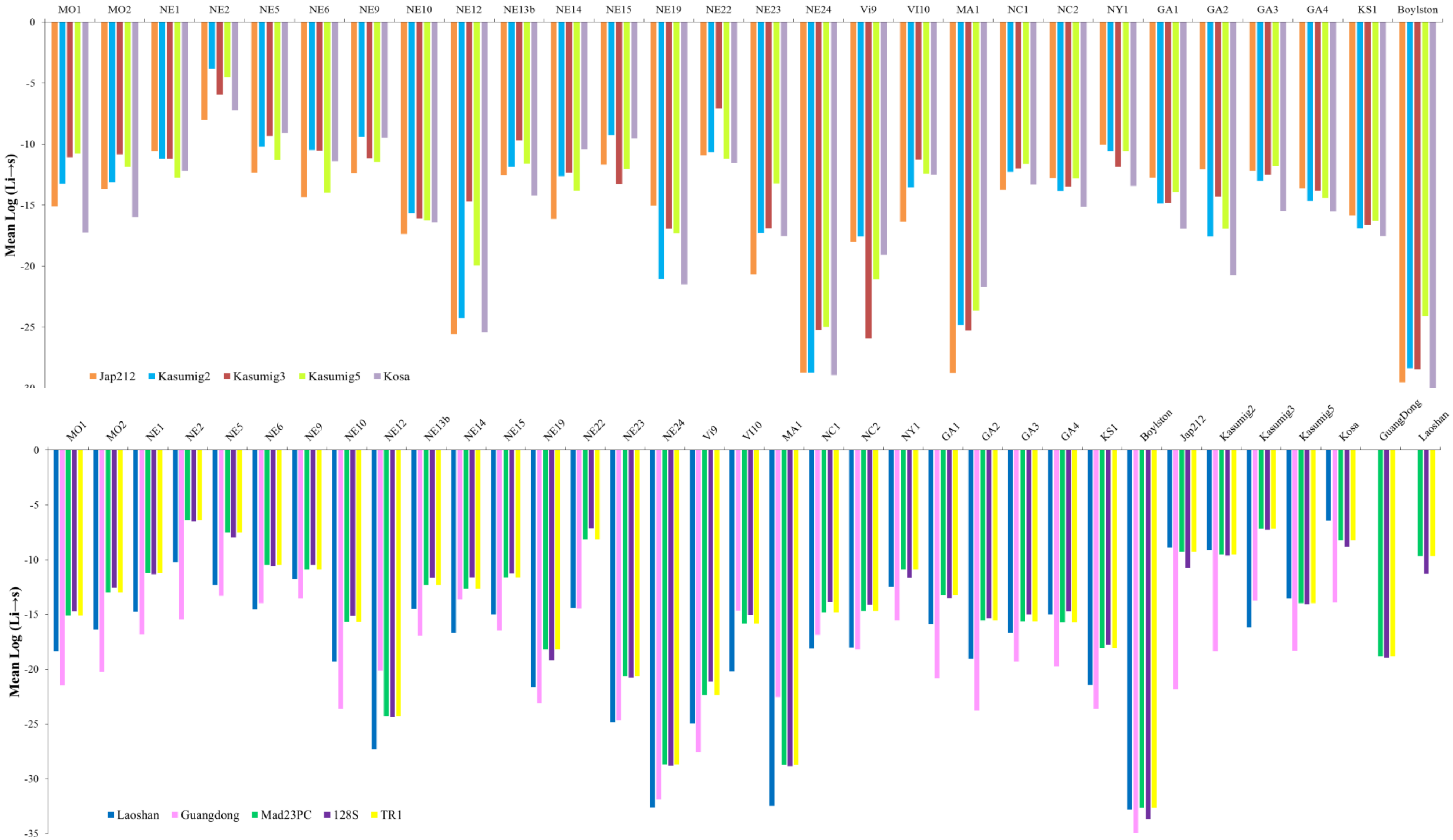
